# Advancing membrane-associated protein docking with improved sampling and scoring in Rosetta

**DOI:** 10.1101/2024.07.09.602802

**Authors:** Rituparna Samanta, Ameya Harmalkar, Priyamvada Prathima, Jeffrey J. Gray

## Abstract

The oligomerization of protein macromolecules on cell membranes plays a fundamental role in regulating cellular function. From modulating signal transduction to directing immune response, membrane proteins (MPs) play a crucial role in biological processes and are often the target of many pharmaceutical drugs. Despite their biological relevance, the challenges in experimental determination have hampered the structural availability of membrane proteins and their complexes. Computational docking provides a promising alternative to model membrane protein complex structures. Here, we present Rosetta-MPDock, a flexible transmembrane (TM) protein docking protocol that captures binding-induced conformational changes. Rosetta-MPDock samples large conformational ensembles of flexible monomers and docks them within an implicit membrane environment. We benchmarked this method on 29 TM-protein complexes of variable backbone flexibility. These complexes are classified based on the root-mean-square deviation between the unbound and bound states (RMSD_UB_) as: rigid (RMSD_UB_ *<*1.2 Å), moderately-flexible (RMSD_UB_ ∈ [1.2, 2.2) Å), and flexible targets (RMSD_UB_ > 2.2 Å). In a local docking scenario, i.e. with membrane protein partners starting ≈10 Å apart embedded in the membrane in their unbound conformations, Rosetta-MPDock successfully predicts the correct interface (success defined as achieving 3 near-native structures in the 5 top-ranked models) for 67% moderately flexible targets and 60% of the highly flexible targets, a substantial improvement from the existing membrane protein docking methods. Further, by integrating AlphaFold2-multimer for structure determination and using Rosetta-MPDock for docking and refinement, we demonstrate improved success rates over the benchmark targets from 64% to 73%. Rosetta-MPDock advances the capabilities for membrane protein complex structure prediction and modeling to tackle key biological questions and elucidate functional mechanisms in the membrane environment. The benchmark set and the code is available for public use at github.com/Graylab/MPDock.

## 1. Introduction

Protein-protein interactions play a pivotal role in biological signaling networks. Elucidating these signaling networks can provide insights into protein function and aid in engineering new therapeutics and de novo protein interfaces. Over the past few years, there have been dramatic advances in protein structure prediction and design(AlphaFold2, ^1^ RFDiffusion,^2^ and Chroma^3^ to name a few); however, most of these advances are biased towards soluble proteins owing to the higher representation of soluble proteins in the Protein Data Bank (PDB).^4^ Membrane protein interactions, *i.e*., interactions between proteins engulfed within lipid bilayers, are one such avenue that is under-studied; with these interactions performing essential life processes ranging from motility and endocytosis, to signaling and sensory responses. The oligomerization of membrane proteins in their native cellular environment plays a fundamental role in the regulation of cellular functions, and their malfunction contributes to a plethora of diseases such as cancer, vascular anomalies, and skeletal syndromes. 5–8 This role has resulted in a major fraction of pharmaceuticals (87% of biologics and 81% of small-molecule drugs) targeting membrane proteins even though membrane proteins span only 30% of all existing natural proteins.^9^ Despite the interest in membrane protein interactions, experimentally determining the precise oligomeric states of membrane proteins remains a challenging problem owing to the heterogenous membrane environment.

Conventionally, membrane protein oligomeric states are characterized in cells or on membrane-mimetic platforms.^10^ While cell-based methods preserve the native cell environment, they often lack high resolution.^11,12^ Conversely, membrane-mimetic platforms offer high molecular resolution but do not replicate the native cell environment^13,14^. The presence of a non-uniform, biphasic membrane layer poses a significant limitation for efficient protein extraction, solubilization, stabilization, and eventually, generation of diffracting crystals or clear cryo-EM grids, hampering structure prediction.^15^ Owing to these challenges, MPs represent less than 3% of all protein structures in the protein data bank (PDB), with MP complexes being even scarcer.^16,17^ When experimental approaches are infeasible, computational modeling tools may address some of these challenges to model MP complexes and protein-protein interactions.

Physics-based computational methods for modeling protein complex structures use a sampling routine and an energy function to approximate the thermodynamics of interactions. On the one hand, the constraints on the search space imposed by the lipid bilayer facilitate docking; on the other hand, the lipid bilayer in tandem with the solvent creates a biphasic environment that complicates modeling. Hence, in spite of several advanced protein docking protocols being available for soluble protein docking, there is a dearth of protocols for membrane protein docking. Conventionally, soluble protein docking protocols are extended for membrane protein docking while rescoring with a membrane-specific energy function. For instance, rigid-body docking algorithms such as DOCK/PIPER^18^ and Memdock^19^ rescore structures using membrane transfer energies in a lipid biphasic environment, but do not consider the membrane during the sampling. Recently, Rudden and Degiacomi developed a membrane docking protocol, Jabberdock^20^, that uses all-atom molecular dynamics to dock proteins while capturing protein backbone motion. Jabberdock first equilibrates the monomers in an explicit membrane environment and then extracts their volumetric mapping to maximize their shape complementarity. On an unbound dataset of 20 *α*-helical complexes of variable flexibility, Jabberdock was successful (*i.e*., yielding at least one acceptable model or better among its top 10 candidates) in 75% of cases (100% for flexible targets). However, the conformational changes sampled are limited by the MD time scale, and the volumetric mapping is computationally expensive (3.5 days on a GPU). Alternatively, to circumvent the limitations of length and timescales with explicit membrane models, Alford *et al*. demonstrated the use of implicit models that represent the membrane as a continuum.^15^ In exchange for an approximate bilayer representation, implicit models offer a 50 − 100 fold sampling speed-up. Implicit membrane models overcome the lipid layer and solvent complexity while maintaining atomic-level details for the molecule of interest. Proof-of-concept work on Rosetta-MPDock^15^ showed this speed-up for rigid-body docking within a membrane-based scoring scheme and has found successful high-ranking poses in three out of five rigid benchmark targets. In that study however, conformational changes were not allowed.

Sampling backbone flexibility upon association has persisted as a long-standing problem even in soluble proteins; evident by limited success rates in capturing flexible proteins in blind structure prediction challenges.^21^ Despite the advent of AlphaFold2 and its breakthrough performance in predicting accurate protein structures, AlphaFold2 (particularly AlphaFold-multimer) predicts only up to 43% of protein complexes accurately. Additionally, AlphaFold2 is found to be less reliable for membrane protein structure prediction.^22^ To address the limitations in flexible membrane protein docking and better sample membrane protein interactions, we present here an update to Rosetta-MPDock that captures binding-induced conformational changes. Rosetta-MPDock mimics the conformer selection mechanism of protein binding by docking large conformational ensembles of membrane protein partners within an implicit membrane environment. Further, we also combined AlphaFold-multimer with Rosetta-MPDock to predict better membrane protein interfaces. This approach is inspired by the improved accuracy that we recently achieved by docking soluble proteins while combining physics and deep-learning based methods.^23^

Here, we first present a curated dataset of 29 trans-membrane protein complexes with variable flexibility that can serve as a benchmark set for validating the performance of membrane protein docking. Next, we demonstrate the performance of Rosetta-MPDock and test whether flexibility improves MP complex structure prediction. Finally, we assess whether AlphaFold-multimer predictions can be used in conjunction with Rosetta-MPDock to predict models with higher recovery of native-like interfaces.

## Results

### Benchmark assembly and method overview

#### Benchmark

To develop and assess computational modeling algorithms, it is crucial to first curate benchmarking datasets. For protein-protein docking, an ideal benchmark set would constitute both bound and unbound conformations of protein partners forming the complex.^24^ One such example is the Docking Benchmark Set (DB 5.5) for soluble protein complexes, which is widely used for evaluating docking performance.^24^ However, for TM protein complexes, the difficulty in experimental characterization has led to the scarcity of both bound and unbound conformations for membrane protein docking.^25^ Prior benchmarks by Almeida *et al*.,^25^ Roel-Torris *et al*.,^26^ and Rudden and Degiacomi^20^ have categorized membrane proteins with respect to their secondary structures (*α*-helical and *β*-sheets), interface locations (cytosolic, TM domain, between TM domains), and their conformational states (bound and unbound). Here, we build on these prior benchmarks to curate a larger, comprehensive dataset of Protein Data Bank (PDB) structures with 29 TM protein complexes and their corresponding unbound conformations. **Table 1** includes each protein target highlighted by its stoichiometry and the extent of flexibility as determined by the unbound-to-bound interface RMSD_UB_ (iRMS). We classified the benchmark set based on the target specifications defined by CAPRI (Critical Assessment of PRedicted Interactions). The current benchmark set comprises 10 moderate to highly flexible targets, encompassing a broad range of interface sizes, sequence lengths. These cleaned and renumbered structures of both unbound and bound conformations are deposited at github.com/Graylab/MPDock to facilitate reproducibility, analysis, and evaluation of alternative membrane modeling tools.

**Table 1.**
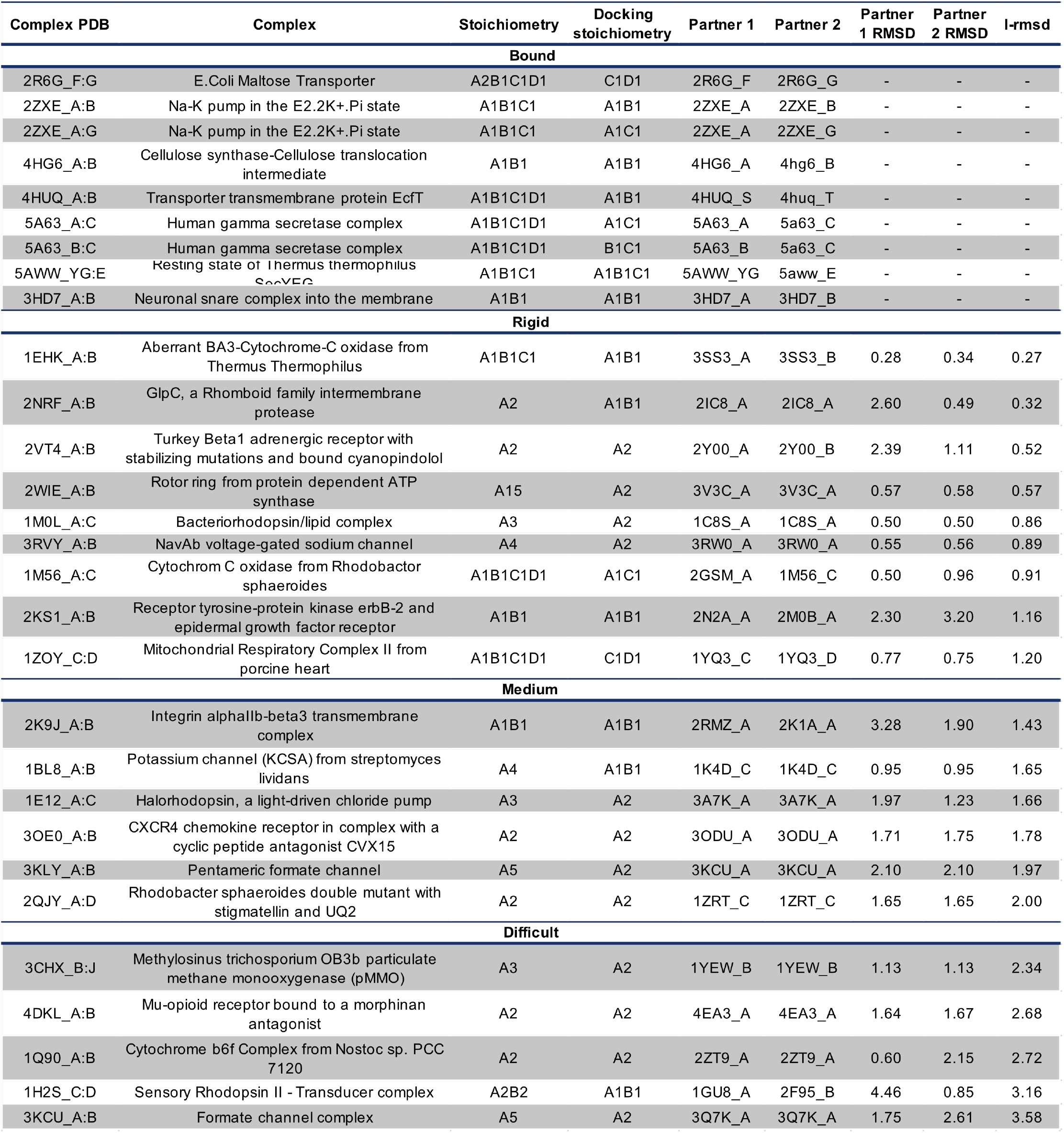
Membrane protein benchmark targets. Benchmark targets organized by flexibility categories: bound (no unbound partners available in the PDB); rigid (RMSD_UB_ < 1.2 Å); moderately-flexible (RMSD_UB_ ∈ [1.2, 2.2) Å); and flexible targets (RMSD_UB_ > 2.2 Å).

#### Rosetta-MPDoc

**Figure 1** illustrates the Rosetta-MPDock protocol with its rigid and ensemble docking versions. Prior work with RosettaMP integrated the membrane-specific environment in Rosetta.^15,27^ The construction of the membrane environment is described in detail by Alford *et al*. and Leman *et al*. respectively and illustrated in **Figure 1.1**.

**Fig. 1.**
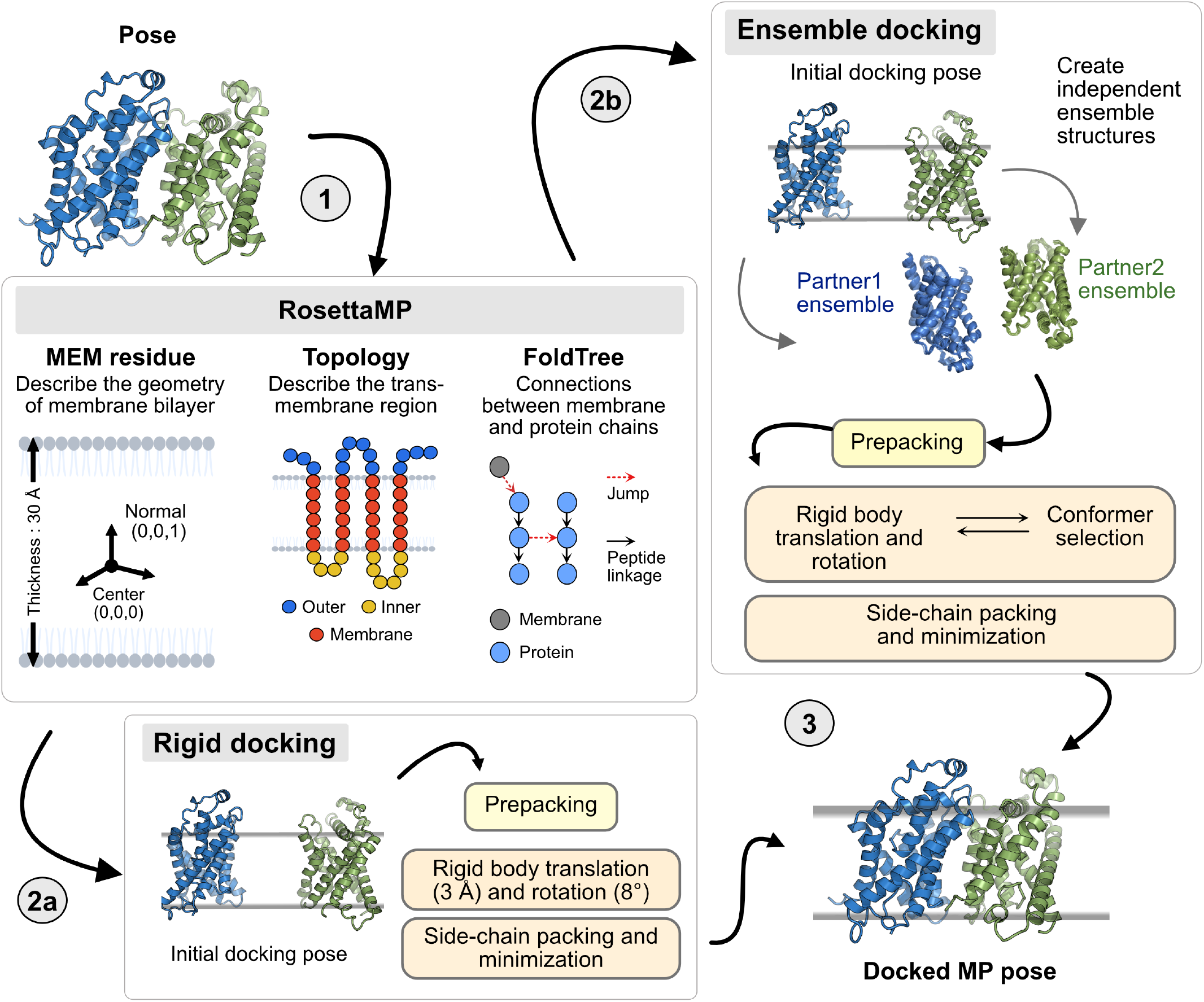
Overview of membrane protein docking protocol. *Panel 1:* RosettaMP architecture: The membrane bilayer is represented using three components namely: MEM residue that describes the geometry of the membrane bilayer; a topology object that stores the transmembrane region information; and a FoldTree object that defines the jump edges to establish the connection between the membrane residue and the protein. *Panel 2a:* Rigid docking protocol with Rosetta-MPDock*Panel 2b:* Ensemble docking protocol with Rosetta-MPDock that involves a conformer-selection approach over an ensemble of pre-generate backbone conformations of the protein partners within the membrane environment. *Panel 3:* A representation of the final docked membrane protein structure that could be obtained from either of the two protocol schemes.

In this work, we use this membrane environment for rigid backbone and flexible backbone protein docking. Rosetta-MPDock performs rigid-body docking by orienting the protein partners in the membrane (as determined by their membrane span files) followed by Monte Carlo moves, i.e. translational and rotational Gaussian perturbations of 3 Å and 8^°^, side-chain packing and relaxation (**Figure 1.2a**). To incorporate conformational changes, we use the conformer selection approach described in RosettaDock 4.0^28^ for soluble proteins. First, structural ensembles for membrane proteins (100 structures for each protein partner) are constructed by Rosetta Relax, Backrub, and Normal Mode Analysis (NMA) while proteins are embedded in a membrane bilayer. Backbone swaps from the ensemble are performed during docking and docked structures are packed and relaxed, then ranked based on their interface scores (i.e. binding energies) to obtain a docked membrane protein complex structure (**Figure 1.2b**). Details are elaborated in **Methods**.

**Fig. 2.**
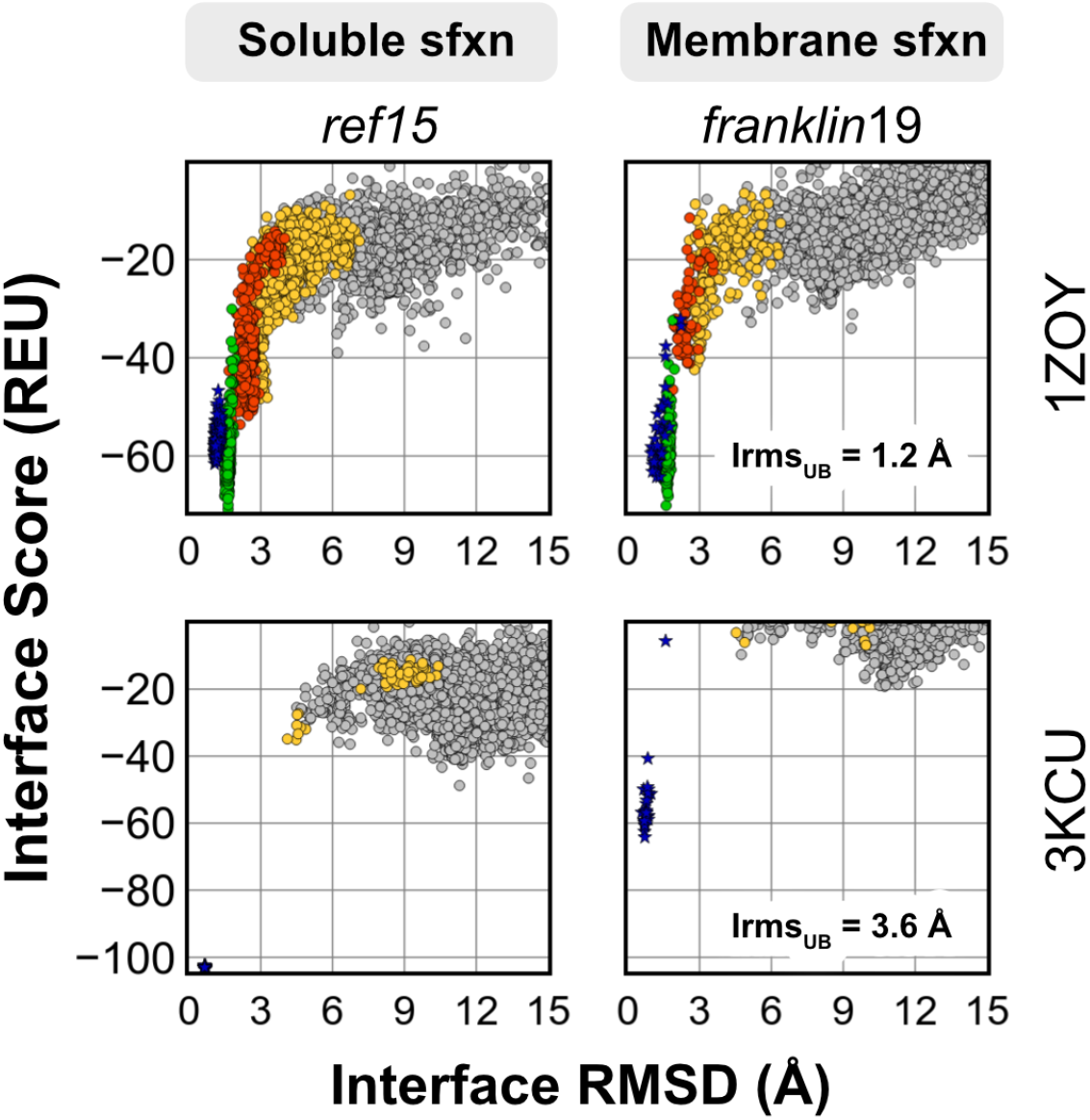
Rigid-body docking energy funnels for protein targets 1ZOY (mitochondrial respiratory complex II, RMSD_*UB*_ = 1.20 Å) and 3KCU (Portable formate transporter, RMSD_*UB*_ = 3.56 Å). Plots show the interface score (REU) vs all-atom C*α* rmsd (Å). Blue stars denote the refined native structures; green, high quality; red, moderate quality; yellow, acceptable quality; gray, incorrect)

### TM-rigid body docking samples high-quality decoys for rigid targets

As a baseline, we first present the performance of two benchmark targets with the Rosetta-MPDock rigid-body protocol for two MP targets a mitochondrial respiratory complex II from porcine heart (1ZOY),1.2 Å RMSD_UB_^29^ and formate channel (3KCU), 3.58 Å RMSD_UB_^30^. **Figure 2** shows the interface score versus the interface RMSD with respect to native for a local docking scenario (protein partners moved 10 Å apart) for two targets across the two scorefunctions. The bound crystal structure is also relaxed to obtain near-native energies (blue stars in Figure 2). For the rigid target 1ZOY, Rosetta-MPDock captures CAPRI high-quality targets (green), and the sampled structures and scores retrace those of the refined near-natives. This is a successful docking scenario. On the other hand, for a flexible target, 3KCU, the performance is underwhelming, with no decoy sampled within 3 Å iRMSD with either scorefunctions. This demonstrates a sampling failure for target 3KCU. This trend is also observed over other medium and highly flexible targets; only 2 out of 11 (18%) medium/highly flexible targets have near-native decoys as opposed to 4 out of 9 (44%) rigid targets (Supplementary Fig. S3-4). While rigid and bound targets are docked with higher accuracy (success rate 56% for 9 targets), the accuracy of flexible targets is hampered despite sampling in the native-like binding region. These results suggest a need to incorporating backbone motions to capture binding-induced conformational changes within membrane-associated protein assemblies.

Next, we compare the discrimination ability of scorefunctions, `ref2015`^31^ (Rosetta energy function for soluble proteins) and `franklin2019`^32^ (Rosetta energy function for membrane proteins). Comparing between the soluble and membrane protein scorefunctions (column-wise panels), we were surprised to observe hardly any improvement in native structure discrimination with the membrane scorefunction. Even though the membrane environment energy terms drive sampling, the high-resolution discrimination at the interface is driven by van der Waals and side-chain packing energy terms, similar to observations in prior work from Alford *et al*.^32^ and Mravic *et al*.^33^

### Ensembles capture binding-induced conformational changes and improve docking performance on flexible targets

To incorporate diverse backbones in membrane protein docking, we developed ensemble docking within Rosetta-MPDock. The ensemble stage in Rosetta-MPDock (**Figure 1**, right panel) draws on the existing conformer-selection functionality of RosettaDock4,^28^ and adapts it for membrane proteins. Conformer-selection^34^ models for protein interactions obey a statistical mechanical view of protein binding; with unbound states of protein partners existing in an ensemble of low-energy conformations, among which the bound conformations are selected during protein association. We implement this strategy by pre-generating an ensemble of conformations of the individual protein partners to use as inputs for docking. While docking, the ligand (smaller protein partner) and the receptor (larger protein partner) undergo rigid body moves coupled with backbone swaps from the pre-generated ensembles. We adapted this strategy for Rosetta-MPDock by implementing the membrane environment for both pre-generating ensembles and making docking moves. By including this backbone diversity, we tested whether we could obtain better near-native sampling for flexible targets.

To demonstrate the performance of ensemble docking vs rigid docking, we compare the docking metrics for the same flexible target 3KCU. **Figure 3A** plots both the interface score (top) and the fraction of native-like contacts made by the interface residues of the sampled decoys with respect to native (bottom) as a function of the interface RMSD. Ensemble docking shows better sampling, as evident from the lower energy decoys sampled within near-native RMSDs and higher *f* _nat_ scores (Figure 3A). This observation supports our hypothesis that backbone sampling allows capturing native-like binding interfaces for flexible targets with considerable conformational change. **Figure 3B** illustrates the best-sampled decoy structure superimposed over the native, highlighting the correct binding orientation in the membrane bilayer being sampled.

**Fig. 3.**
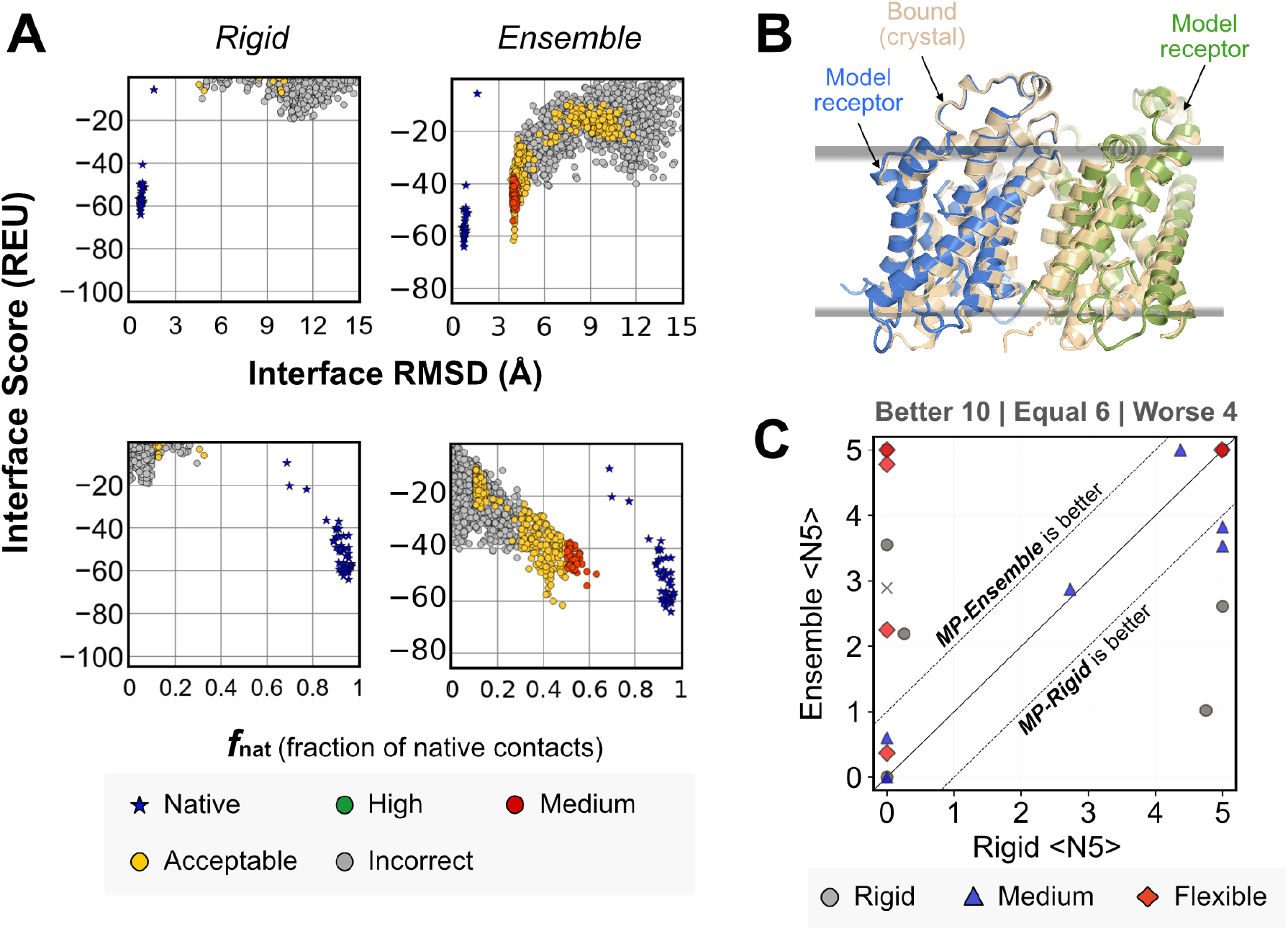
Ensemble-MPDock improves docking performance on flexible targets. (A) Interface Score (REU) vs Interface RMSD (Å) (top), and fraction of native-like contacts (bottom) for target 3KCU. (B) Best sampled decoy for 3KCU (portable formate transporter, RMSD_*UB*_ = 3.56 Å). (C) Comparison of ⟨N5⟩ values after full protocol for Rosetta-MPDock *rigid* and *ensemble* cases respectively. Dashed lines highlight the region in which the two protocols differ significantly, i.e. by more than one point in their ⟨N5⟩ values. Different symbols correspond to each target’s difficulty category (circle: rigid; triangle: medium; diamond: flexible). Points above the solid line represent better performance with *franklin*19 scorefunction, while points below the line represent better performance with mp15 scorefunction.

Next, to compare the two scorefunctions, we measured the number of near-native decoys in the top 5 structures (⟨N5⟩) for the full benchmark set of 29 targets. A near-native structure is considered to be a success if it is a decoy with a CAPRI rank of acceptable or higher. The protein target is considered a docking success if three of the top five scored structures are near-native, accessed with bootstrapped sampling (⟨N5⟩≥3). **Figure 3C** compares the ⟨N5⟩ scores of rigid docking and ensemble docking with the dashed lines signifying the region of little difference. Targets in the upper half indicate that the ensemble docking performs better, whereas those in the lower half indicate that rigid docking performs better. Almost all flexible targets (red diamonds) exhibit equal or better performance with ensemble docking. However, for medium targets (blue triangles), ensemble docking often reduces the performance. The docking funnel plots (Supplementary S3-S4) show that although lower RMSD structures are sampled, some docking trajectories led to false positive minima, suggesting a need to improve the energy function. The false positive minima could also arise from backbone motion in regions of the protein that do not move in reality, resulting in an unrealistic backbone conformation that seems to fit better *in silico*. Overall, the improvement by `franklin19` scorefunction is modest. `Franklin2019` focuses on the hydrophobic interaction between the proteins and the membrane bilayer however, it misses the electrostatic interaction (Supplementary figures S1-S6). Recently, we developed a new energy function, `franklin23`, to add the electrostatic effect of the phospholipid layer and variable dielectric constant in the membrane bilayer. A comparison of interface rmsd and the fraction of native contacts by `franklin23` shows similar or slightly better results in comparison to `franklin2019` as shown in Supplementary Figures S7 and S8. scorefunctions and their details are discussed in SI section 1. Irrespective of functions, ensemble docking improves docking for flexible targets over conventional rigid body docking (Supplementary Fig. S5-6).

### MPDock efficiently refines AlphaFold predictions and recapitulates native-like contacts

Deep-learning approaches such as AlphaFold2 and RoseTTAFold have enabled highly accurate three-dimensional structure prediction. Further, AlphaFold-multimer (AFm) has improved the structure prediction of protein complexes, however flexible protein complexes and transmembrane proteins are still a challenge.^23,35,36^ Here, we assess the performance of AFm for membrane protein assemblies on the benchmark set. Note that most of the targets were deposited in the Protein Data Bank (PDB) before AFm’s training date, so performance on novel structures may be worse. We evaluate whether refining AFm predicted structures with Rosetta-MPDock (AFm+Rosetta MPDock) can improve performance. **Figure 4A** shows the RMSDs for medium and flexible targets of the benchmark set across different docking protocols (starting from the unbound conformers) compared to the AFm predicted structures. We compare the *C*_*α*_ RMSDs of the protein complexes obtained from prediction tools (AFm, JabberDock, Rosetta MPDock, and AFm+Rosetta MPDock) with the experimental structures. AFm results are highlighted as a red cross. In comparison to AFm and JabberDock, AFm+Rosetta MPDock (rigid and ensemble) captures lower RMSD structures. For cases of interface rmsd over5 Å, for instance, targets 3CHX, 4DKL, and 1Q90, the higher interface rmsd may be explained by poor prediction of protein partner structures, i.e., if individual protein partners were themselves predicted incorrectly, MPDock protocol fails to dock them successfully. Therefore, a major limitation in utilizing docking protocols over structure prediction tools is that the accuracy of docking would depend upon the prediction accuracy of protein partners.

**Fig. 4.**
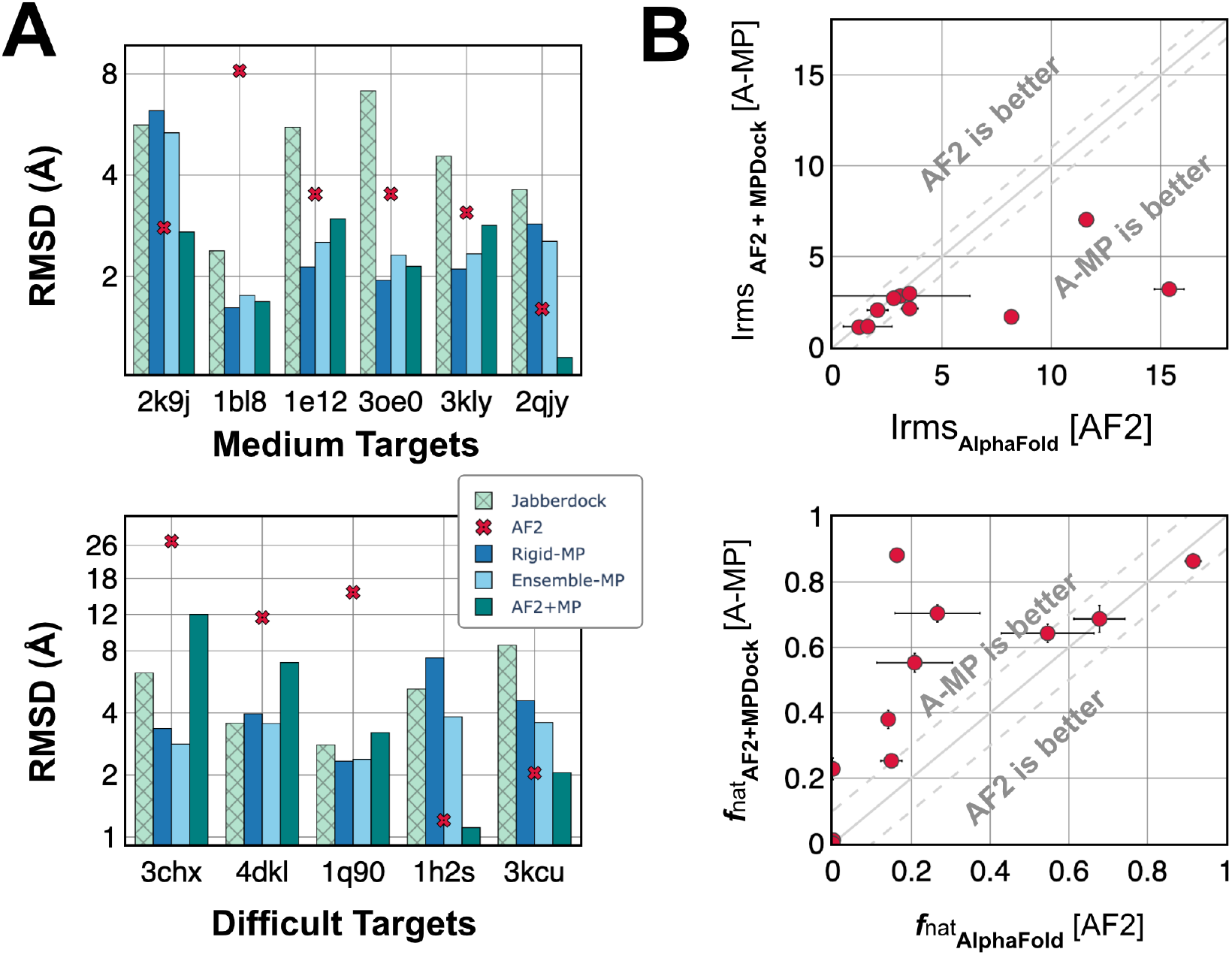
Performance of MPDock with AlphaFold2 predicted structures. (A) Interface RMSD (*on top*) and fraction of native-like contacts (f_nat_) for Rosetta-MPDock (ensemble and rigid docking starting from unbound monomers) and AFm+RosettaMP dock (ensemble and rigid docking starting from Alphafold2 predicted monomers). Performance is indicated by lower Irmsd ad higher f_nat_). (B) RMSD of the predicted protein monomer (*shaded*) or protein complex (*blank*) structure relative to the native/bound crystal structure for moderately flexible/medium (*top*) and difficult targets (*bottom*).

To obtain a head-to-head comparison between AFm and AFm+MPDock (ensemble), we compare the interface RMSDs and *f* _nat_ of top-5 structures from respective methods (**Figure 4B**). Alphafold2 captures near-native structures in a few cases, but we observe that in almost all the cases, Rosetta MPDock refinement improved Alphafold2 predictions to capture near-native structures, evident from the lower interface RMSD (Irms) and a higher fraction of native-like contacts (*f*_nat_) for Rosetta MPDock. Thus, refinement and docking with a physics-based scorefunction that accounts for the membrane environment can generate better membrane protein assemblies.

## Discussion

Despite their significant importance as pharmaceutical drug targets, structure determination is notoriously difficult for membrane proteins. In this work, we developed, benchmarked, and evaluated a docking pipeline that accommodates the membrane environment and enables flexible backbone protein-protein docking. We built on the foundations of RosettaMembrane modeling tools to create a modular framework for membrane protein docking with backbone flexibility. Rosetta-MPDock combines the features of the membrane environment (membrane topology, span, and geometry) with docking features and a conformer-selection mechanism to provide a membrane protein docking algorithm. Further, by incorporating Alphafold2 modeled structures and assessing them in energy functions suitable for a membrane-specific environment, we demonstrate an ability to sample better docked models. The results on a membrane protein benchmark of 29 targets improve membrane protein structure determination and lay the groundwork for answering underlying questions in biology involving trans-membrane proteins.

The membrane protein docking benchmark that we curated, is to the best of our knowledge, the most comprehensive database of transmembrane protein structures with known bound and unbound forms. We further demonstrated the utility of a flexible backbone protocol over conventional rigid-body docking approaches in sampling moderately-flexible and flexible targets. By incorporating diverse backbones generated from different ensemble generation protocols along with an improved membrane energy function, Rosetta-MPDock can effectively identify near-native interfaces. This is reflected by a boost in docking performance relative to alternative state-of-the-art docking methods (e.g. HADDOCK, JabberDock) as Rosetta-MPDock successfully docks 67% of moderately flexible targets and 60% of flexible targets.

One of the limiting factors in conformer-selection methods has been the difficulty of ensemble-generation methods in capturing native-like structures. With the advent of Alphafold2 (and recently AlphaFold3^37^), there is an opportunity to leverage its structural predictions to diversify conformational ensembles and provide plausible backbones for protein docking. We demonstrate this by coupling AlphaFold2 predictions with Rosetta-MPDock. In cases where AlphaFold2 predicts unbound protein partners with high accuracy, Rosetta-MPDock refines on those inputs to create CAPRI-acceptable or better models. We have previously shown that augmenting AlphaFold2 with physics-based sampling strategies has demonstrated potential for soluble protein docking and antibody-antigen targets.^23^ Our results here extend these observations for membrane proteins and show that physics fused with deep learning structure prediction tools can guide better sampling in the relatively difficult challenge of sampling membrane protein conformations. We anticipate that the availability of the benchmark and the modeling tools will make membrane protein modeling accessible to the broad scientific community and enable better design of this exquisite class of biomolecules.

## Methods

### Dataset Curation

We built on prior benchmarks^20,25^ and curated a consolidated set with 29 TM proteins and their unbound conformations. We classified these complexes based on their extent of flexibility, (unbound-to-bound root-mean-square-deviation for interface residues, RMSD_unbound-bound_) into the following categories: bound (with no unbound conformations available); rigid; medium; and difficult. The curated benchmark set features 9 bound targets, 9 rigid targets, 6 medium targets, and 5 difficult targets (**Table 1**).

### Energy functions

We tested *franklin*19^32^, the current standard for membrane protein modeling in Rosetta; and three different membrane energy functions along with one soluble protein energy function in our benchmarking analysis. The membrane energy functions were membrane protein framework 2015 (`MP15`)^15^, franklin2019 (`franklin19`)^32^ and franklin2023 (`franklin23`,^16^ new energy function *in Supplementary*); with ref2015 (*ref* 15)^15^ as the soluble energy function. Further details about individual energy functions are described in the supplement. All the energy functions correspond to Rosetta’s all-atom mode and have been benchmarked on experimental metrics such as tilt angle, stability, and design.^17^ We use the motif dock score (MDS) energy function for the low-resolution phase in Rosetta MPdocking protocols due to the lack of a membrane-based low-resolution version of `franklin19`. MDS relies on a pre-calculated residue pair energy that resembles ref15 energies mapped onto backbone coordinates; however, it lacks the membrane context.

### Rosetta MPDock protocol

#### Rigid docking

Rosetta MPDock^15^ is an extension of the conventional RosettaDock protocol to incorporate the complexities of modeling membrane proteins. Rosetta MPDock protocol transforms the input pose to the membrane environment, pre-packs the input structure (optimizing rotameric conformations for side chains) and then engages in docks within the lipid membrane with rigid-body rotations and translations performed in 2D cartesian space (*x, y* coordinate space as the *z* coordinate is constant owing to membrane-depth). The lipid membrane is fixed throughout the sampling procedure, and each sampled conformation is scored with a membrane-specific scorefunction. The details of the protocols are in Alford and Leman *et. al*.^15^

#### Ensemble docking

Building over the Rosetta MPDock rigid-body docking protocol, ensemble docking incorporates diverse backbones to mimic conformer selection in docking. Following the transformation of the `Pose` object into the membrane environment, the ensemble docking protocol performs three steps: (1) ensemble generation to diversify the protein backbone, (2) the pre-packing to refine the side chains and create a starting structure, and (3) protein-protein docking in the membrane bilayer. In the **ensemble generation** step, to generate diversity in backbone conformations for the proteins, we used three conformer generation methods: perturbation of the backbones along the normal modes by 1 Å^38^ using RosettaScripts^39^ refinement using the Relax protocol in Rosetta,^40^ and backbone variation using the Rosetta Backrub protocol.^41^ Complete command lines are provided in the Supplementary Method. We have used 40 Backrub conformers, 30 normal mode conformers, and 30 relax conformers to comprise an ensemble of 100 conformers. Similar to the rigid docking, in the **pre-packing step**, the side chains of the ensembles of the unbound structures (keeping their membrane embedding constant) are repacked using rotamer trials. Next, the **docking step** uses a Monte Carlo plus a minimization algorithm^42^ consisting of a low-resolution stage simulating conformer selection and a high-resolution stage simulating induced fit. The low-resolution stage includes rotating and translating the ligand around the receptor coupled with swapping of the pre-generated backbone conformations using Adaptive Conformer Selection.^28^ In the high-resolution stage, the side chains are reintroduced to the putative encounter complex, and those at the interface are packed for tight binding. At all steps, the membrane is kept fixed.

## Data Availability

The source code for docking, with interface-tests, global and local docking examples and directed induced-fit, is available at rosettacommons.org, including scripts and tutorials. The benchmark and other utility scripts are available at github.com/Graylab/MPDock.

## Conflicts of Interest

JJG is an unpaid board member (director) of the Rosetta Commons. Under institutional participation agreements between the University of Washington, acting on behalf of the Rosetta Commons, Johns Hopkins University may be entitled to a portion of the revenue received on licensing Rosetta software, including some methods described in this manuscript. JJG has a fiduciary role in Levitate Bio LLC. Levitate Bio LLC distributes the Rosetta software, which may include methods described in this paper. Janssen Research Development, LLC has licensed Rosetta and PyRosetta software from University of Washington who manages the licensing on behalf of the Rosetta Commons. JJG provides paid consulting services to Janssen Research Development, LLC. JJG has a financial interest in Cyrus Biotechnology. These arrangements have been reviewed and approved by Johns Hopkins University in accordance with its conflict-of-interest policies.

## ACKNOWLEDGMENTS

This work was supported by the National Institute of Health through grants R35-GM141881 and R01-GM078221. Computational resources were provided by the Extreme Science and Engineering Discovery Environment (XSEDE) and Advanced Research Computing at Hopkins (ARCH). We appreciate Hope Woods for reviewing our code and Sergey Lyskov for helping us with the server and demos.

