## Supplementary File for "Advancing membrane-associated protein docking with improved sampling and scoring in Rosetta"

#### An expanded benchmark for flexible backbone transmembrane docking

Figs. S1 to S10
Tables S1 to S4
SI References

### 1. Supplementary Results

#### Energy function details.

MP15 energy function uses different representations for low and high-resolution terms. At low resolution, the membrane comprises hydrophobic core, interface, and solvent regions.(? ) In contrast, the full atom high-resolution mode uses the Lazaridis membrane model (? ) to describe a continuous dielectric gradient from the hydrophobic core to the solvent. The low-resolution energy terms were based on environment, pair, and density terms of 28 TM helical proteins. On the other hand, MP15 uses transfer energy from water to octanol to represent the transfer energy for residues and a few other knowledge-based environment terms. The functional form of each of the terms is given in Table B of the supplementary section of Alford and Leman et al.(? ) *franklin*19 uses the all-atom energy function `ref15` for soluble proteins and added the transfer energy of residues in the membrane based on the Moon and Fleming (MF) scale measuring the transfer energy of residues in the phospholipid bilayer.(? ) `ref15` computes macromolecular energies through a linear combination of terms for van der Waals, solvation, electrostatics, hydrogen bonding, backbone, and side-chain interactions.(1) *franklin*19 included MP pores and features of different phospholipids in the model. Recently, we developed a new energy function, namely *franklin23*, to add the electrostatic effect of the phospholipid layer and variable dielectric constant in the membrane bilayer. This energy function is built over *franklin19* and in this work, we assess the performance over 2 protein targets ( a rigid target 1ZOY and a flexible target 3KCU) from the benchmark. A comparison of i-rmsd and fraction of native contacts by *franklin23* shows similar or slightly better results in comparison to *franklin2019* as shown in Supplementary Fig. S7 and S8. The functional forms of *franklin*19 and *franklin*23 are:

$$\textit{franklin}23 = \text{ref15} + w_{w,l}\Delta G_{w,l} + w_{\text{lipid}}\Delta G_{\text{lipid}} + w_{\text{dielec}}\Delta E^{\text{dielectric}}_{\text{elec},ij} \quad [1]$$

*where* $\Delta G_{w,l}$ presenting the transfer energy from water to bilayer(? ); $\Delta G_{\text{lipid}}$ and $\Delta E^{\text{dielectric}}_{\text{elec},ij}$ are the energies due to lipid head group and the membrane dielectric constant respectively.(? )

We can recover the energy function *franklin*19 by setting $w_{w,l} = 0.5$, $w_{\text{lipid}} = 0$ and $w_{\text{dielec}} = 0$ and *franklin*23 by assigning $w_{w,l} = 1, w_{\text{lipid}} = 0.128$ and $w_{\text{dielec}} = 0.01$. Both these energy functions are all-atom and have been benchmarked on experimental measures such as tilt angle, stability, and design.(? ? ) Due to the lack of a low-resolution version of *franklin*19 and *franklin*23 energy functions, we have used the motif dock score (MDS) low-resolution energy function. MDS relies on a pre-calculated residue pair energy that resembles `ref15` energies mapped onto backbone coordinates, however, lacks the membrane context. Due to the benefits and drawbacks of all score functions, we decided to test them all. Even though the performance of the high-resolution MP15 score function has been compared with Franklin energy functions before, here we take the opportunity to evaluate the combination of low- and high-score functions on a larger docking dataset.

**Table S1. Summary of average ⟨N5⟩ values of protein complexes docked using RosettaMP rigid and ensemble docking protocols using MP15 and Franklin2019 (F19) score functions. The average ⟨N5⟩ values for MPDock starting with AF2 structures are shown for rigid docking in AF_MPRigid and for ensemble docking in AF_MPEnsemble. Results for MPDock starting from AF2 structures are scored with F19.**

| Complex PDB | Flexibility | $RMSD_{BU}$ | Ensemble_F19 average N5 | Ensemble_MP15 average N5 | Rigid_F19 average N5 | Rigid_MP15 average N5 | AF_MPRigid average N5 | AF_Mpensemble average N5 |
|---|---|---|---|---|---|---|---|---|
| 1EHK | Rigid | 0.27 | 2.69 | 0.00 | 5.00 | 5.00 | | |
| 2NRF | Rigid | 0.32 | 5.00 | 4.80 | 5.00 | 5.00 | | |
| 2VT4 | Rigid | 0.52 | 3.55 | 4.17 | 0.00 | 0.00 | | |
| 2WIE | Rigid | 0.57 | 5.00 | 5.00 | 5.00 | 5.00 | | |
| 1M0L | Rigid | 0.86 | 0.00 | 0.00 | 0.00 | 1.85 | | |
| 3RVY | Rigid | 0.89 | 0.96 | 1.93 | 4.79 | 5.00 | | |
| 1M56 | Rigid | 0.91 | 5.00 | 4.95 | 0.00 | 0.00 | | |
| 2KS1 | Rigid | 1.16 | 2.22 | 0.00 | 0.24 | 0.00 | | |
| 1ZOY | Rigid | 1.20 | 5.00 | 4.89 | 5.00 | 5.00 | | |
| 2K9J | Medium | 1.43 | 0.00 | 0.00 | 0.00 | 0.00 | 5.00 | 1.00 |
| 1BL8 | Medium | 1.65 | 3.90 | 5.00 | 5.00 | 5.00 | 0.00 | 1.00 |
| 1E12' | Medium | 1.66 | 5.00 | 4.90 | 4.33 | 3.10 | 4.02 | 0.00 |
| 3OE0 | Medium | 1.78 | 2.92 | 0.00 | 2.73 | 3.19 | 0.00 | 0.74 |
| 3KLY | Medium | 1.97 | 3.61 | 3.96 | 5.00 | 5.00 | 5.00 | 1.00 |
| 2QJY | Medium | 2.00 | 0.60 | 0.04 | 0.00 | 3.81 | 2.56 | 1.00 |
| 3CHX | Difficult | 2.34 | 0.35 | 0.80 | 0.00 | 0.00 | 0.00 | 0.00 |
| 4DKL | Difficult | 2.68 | 2.18 | 0.75 | 0.00 | 0.42 | 0.00 | 0.00 |
| 1Q90 | Difficult | 2.72 | 5.00 | 3.73 | 5.00 | 3.97 | 0.00 | 0.00 |
| 1H2S | Difficult | 3.16 | 5.00 | 0.78 | 0.00 | 0.00 | 5.00 | 1.00 |
| 3KCU | Difficult | 3.58 | 4.77 | 1.16 | 0.00 | 0.00 | 5.00 | 1.00 |

**Table S2. Summary of fraction of native-like contacts ($f_{nat}$) values for 10 randomly chosen decoys of protein complexes docked using RosettaMP rigid and ensemble docking protocols with mp15, F19 score functions and JabberDock method. The average $f_{nat}$ values for MPDock starting with AF2 structures are shown for rigid docking in AF_MPRigid and for ensemble docking in AF_MPEnsemble. Results for MPDock starting from AF2 structures are scored with F19.**

| Complex PDB | Flexibility | $RMSD_{BU}$ | Ensemble_F19 fnat N10 | Ensemble_MP15 fnat N10 | Rigid_F19 fnat N10 | Rigid_MP15 fnat N10 | AF_MPRigid fnat N10 | AF_Mpensemble fnat N10 | JabberDock fnat |
|---|---|---|---|---|---|---|---|---|---|
| 1EHK | Rigid | 0.27 | 0.10 | 0.05 | 0.19 | 0.19 | | | 0.28 |
| 2NRF | Rigid | 0.32 | 0.85 | 0.76 | 0.95 | 0.92 | | | 0.41 |
| 2VT4 | Rigid | 0.52 | 0.36 | 0.31 | 0.00 | 0.15 | | | 0.42 |
| 2WIE | Rigid | 0.57 | 0.68 | 0.67 | 0.88 | 0.88 | | | 0.65 |
| 1M0L | Rigid | 0.86 | 0.06 | 0.00 | 0.00 | 0.25 | | | 0.87 |
| 3RVY | Rigid | 0.89 | 0.12 | 0.12 | 0.55 | 0.66 | | | 0.25 |
| 1M56 | Rigid | 0.91 | 0.56 | 0.32 | 0.00 | 0.00 | | | 0.00 |
| 2KS1 | Rigid | 1.16 | 0.18 | 0.08 | 0.09 | 0.05 | | | 0.08 |
| 1ZOY | Rigid | 1.20 | 0.76 | 0.35 | 0.88 | 0.90 | | | 0.49 |
| 2K9J | Medium | 1.43 | 0.00 | 0.00 | 0.00 | 0.00 | 0.91 | 0.69 | 0.79 |
| 1BL8 | Medium | 1.65 | 0.49 | 0.64 | 0.94 | 0.93 | 0.16 | 0.88 | 0.47 |
| 1E12' | Medium | 1.66 | 0.18 | 0.19 | 0.22 | 0.20 | 0.22 | 0.25 | 0.66 |
| 3OE0 | Medium | 1.78 | 0.23 | 0.10 | 0.36 | 0.29 | 0.08 | 0.38 | 0.48 |
| 3KLY | Medium | 1.97 | 0.31 | 0.20 | 0.82 | 0.85 | 0.80 | 0.55 | 0.70 |
| 2QJY | Medium | 2.00 | 0.12 | 0.02 | 0.00 | 0.22 | 0.53 | 0.70 | 0.68 |
| 3CHX | Difficult | 2.34 | 0.06 | 0.07 | 0.01 | 0.00 | 0.00 | 0.00 | 0.95 |
| 4DKL | Difficult | 2.68 | 0.15 | 0.15 | 0.01 | 0.07 | 0.00 | 0.01 | 0.96 |
| 1Q90 | Difficult | 2.72 | 0.43 | 0.15 | 0.41 | 0.31 | 0.01 | 0.23 | 0.86 |
| 1H2S | Difficult | 3.16 | 0.31 | 0.07 | 0.00 | 0.00 | 0.94 | 0.86 | 0.48 |
| 3KCU | Difficult | 3.58 | 0.57 | 0.13 | 0.06 | 0.04 | 0.88 | 0.64 | 0.74 |

**Table S3. Summary of average of interface RMSD (IRMSD) values for 10 randomly chosen decoys of protein complexes docked using RosettaMP rigid and ensemble docking protocols using mp15 and F19 score functions. Average IRMSD values for MPDock starting with AF2 structures are shown for rigid docking in AF_MPRigid and for ensemble docking in AF_MPEnsemble. Results for MPDock starting from AF2 structures are scored with F19.**

| Complex PDB | Flexibility | $RMSD_{BU}$ | Ensemble_F19 average IRMS N10 | Ensemble_MP15 average IRMS N10 | Rigid_F19 average IRMS N10 | Rigid_MP15 average IRMS N10 | AF_MPRigid average IRMS N10 | AF_Mpensemble average IRMS N10 |
|---|---|---|---|---|---|---|---|---|
| 1EHK | Rigid | 0.27 | 5.04 | 5.94 | 4.21 | 4.19 | | |
| 2NRF | Rigid | 0.32 | 1.44 | 2.15 | 1.19 | 1.50 | | |
| 2VT4 | Rigid | 0.52 | 4.06 | 3.99 | 21.43 | 5.87 | | |
| 2WIE | Rigid | 0.57 | 1.70 | 1.65 | 1.25 | 1.25 | | |
| 1M0L | Rigid | 0.86 | 5.93 | 10.14 | 11.87 | 8.78 | | |
| 3RVY | Rigid | 0.89 | 10.44 | 10.57 | 3.22 | 1.72 | | |
| 1M56 | Rigid | 0.91 | 2.11 | 3.82 | 16.09 | 16.63 | | |
| 2KS1 | Rigid | 1.16 | 7.56 | 7.28 | 6.99 | 6.25 | | |
| 1ZOY | Rigid | 1.20 | 2.07 | 3.55 | 1.64 | 1.45 | | |
| 2K9J | Medium | 1.43 | 8.46 | 8.38 | 9.20 | 9.05 | 2.82 | 2.71 |
| 1BL8 | Medium | 1.65 | 3.68 | 2.45 | 1.67 | 1.68 | 8.34 | 1.68 |
| 1E12' | Medium | 1.66 | 4.33 | 4.37 | 4.41 | 4.55 | 4.43 | 2.96 |
| 3OE0 | Medium | 1.78 | 6.98 | 5.84 | 8.06 | 5.05 | 6.71 | 2.14 |
| 3KLY | Medium | 1.97 | 6.06 | 4.07 | 2.14 | 2.10 | 2.90 | 2.83 |
| 2QJY | Medium | 2.00 | 10.56 | 12.00 | 26.21 | 5.29 | 5.68 | 1.15 |
| 3CHX | Difficult | 2.34 | 6.80 | 8.01 | 11.37 | 16.47 | 36.38 | 12.00 |
| 4DKL | Difficult | 2.68 | 6.48 | 4.72 | 16.87 | 6.07 | 14.07 | 7.03 |
| 1Q90 | Difficult | 2.72 | 3.91 | 4.36 | 3.15 | 3.92 | 16.55 | 3.20 |
| 1H2S | Difficult | 3.16 | 4.32 | 6.58 | 10.26 | 10.26 | 1.41 | 1.11 |
| 3KCU | Difficult | 3.58 | 4.46 | 9.80 | 10.98 | 12.92 | 2.03 | 2.05 |

**Table S4. Summary of best interface RMSD (Å) values for decoys of protein complexes docked using RosettaMP rigid and ensemble docking protocols using mp15, F19 score functions, and JabberDock method. The best IRMSD values for MPDock starting with AF2 structures are shown for rigid docking in AF_MPRigid and for ensemble docking in AF_MPEnsemble. Results for MPDock starting from AF2 structures are scored with F19.**

| Complex PDB | Flexibility | $RMSD_{BU}$ | Ensemble_F19 best IRMS | Ensemble_MP15 best IRMS | Rigid_F19 best IRMS | Rigid_MP15 best IRMS | AF_MPRigid best IRMS | AF_Mpensemble best IRMS | JabberDock best IRMS |
|---|---|---|---|---|---|---|---|---|---|
| 1EHK | Rigid | 0.27 | 4.35 | 4.75 | 4.18 | 4.18 | | | 5.25 |
| 2NRF | Rigid | 0.32 | 1.16 | 1.24 | 0.76 | 0.97 | | | 9.88 |
| 2VT4 | Rigid | 0.52 | 1.70 | 2.61 | 1.17 | 2.62 | | | 5.11 |
| 2WIE | Rigid | 0.57 | 1.47 | 1.43 | 1.21 | 1.17 | | | 3.54 |
| 1M0L | Rigid | 0.86 | 2.34 | 2.40 | 2.18 | 1.17 | | | 5.26 |
| 3RVY | Rigid | 0.89 | 1.39 | 2.64 | 1.61 | 1.50 | | | 17.66 |
| 1M56 | Rigid | 0.91 | 1.95 | 2.30 | 12.91 | 13.12 | | | 37.12 |
| 2KS1 | Rigid | 1.16 | 5.31 | 4.90 | 5.26 | 5.14 | | | 13.26 |
| 1ZOY | Rigid | 1.20 | 1.71 | 2.08 | 1.49 | 1.35 | | | 12.57 |
| 2K9J | Medium | 1.43 | 5.31 | 5.98 | 5.61 | 5.81 | 2.57 | 2.66 | 5.63 |
| 1BL8 | Medium | 1.65 | 1.73 | 2.00 | 3.31 | 1.57 | 5.91 | 1.67 | 2.38 |
| 1E12' | Medium | 1.66 | 2.41 | 2.73 | 2.09 | 2.32 | 2.98 | 2.79 | 5.55 |
| 3OE0 | Medium | 1.78 | 2.30 | 2.82 | 1.89 | 2.12 | 2.34 | 2.09 | 7.12 |
| 3KLY | Medium | 1.97 | 2.29 | 2.54 | 2.09 | 2.05 | 2.82 | 2.81 | 4.55 |
| 2QJY | Medium | 2.00 | 2.51 | 3.73 | 2.55 | 2.74 | 0.91 | 1.11 | 3.62 |
| 3CHX | Difficult | 2.34 | 2.79 | 3.04 | 3.28 | 2.98 | 13.93 | 11.17 | 6.25 |
| 4DKL | Difficult | 2.68 | 3.44 | 3.67 | 3.90 | 3.68 | 6.49 | 6.90 | 3.57 |
| 1Q90 | Difficult | 2.72 | 2.34 | 2.73 | 2.32 | 2.27 | 7.28 | 3.08 | 2.80 |
| 1H2S | Difficult | 3.16 | 3.80 | 3.95 | 7.36 | 7.86 | 1.03 | 1.09 | 5.22 |
| 3KCU | Difficult | 3.58 | 3.55 | 4.70 | 4.53 | 5.28 | 1.96 | 2.03 | 8.52 |

#### 2. Supplementary Methods

1. Download membrane-embedded PDB file

   Generate a PDB file where the membrane protein structure is transformed into PDB coordinates (z-axis is membrane normal). This can be done either by downloading the transformed PDB directly from PDBTM website or by uploading a PDB file to PPM server.

2. Clean the PDB file

```
$ /path/to/Rosetta/main/tools/protein_tools/scripts/clean_pdb.py 1AFO_tr.pdb
    ignorechain
```

3. Span File: Generate a span file from PDB (1AF0) structure

```
$ /path/to/Rosetta/main/source/bin/spanfile_from_pdb.linuxgccrelease -database /
   path/to/db -in:file:s example_inputs/1AFO_tr.pdb
```

   This generates three span files in the input folder. For this demo, these files have been moved into the output folder.

```
$ 1AFO_AB.span #spanfile of full PDB
$ 1AFO_ABA.span #spanfile of chain A
$ 1AFO_ABB.span #spanfile of chain B
```

4. Ensemble Generation

   (a) Backrub

```
$ /path/to/Rosetta/main/source/bin/backrub.linuxgccrelease @backrub_flags
@backrub_flags
-in:file:s complex_A.pdb #chainA of the unbound complex
-nstruct 40 #number of desired number of ensembles
-backrub:ntrials 20000 #value of kT for Monte Carlo
-backrub:mc_kt 0.6 #value of kT for Monte Carlo
-out:path:pdb output #name of folder to save pdb
-out:prefix br_ #Prefix for output pdb names
-out:pdb_gz #Compress (gzip) output pdbs
-out:path:score output #name of folder to save score of pdbs
```

   (b) Relax

```
$ /path/to/Rosetta/main/source/bin/relax.linuxgccrelease @relax_flags
@relax_flags
-in:file:s complex_A.pdb #chainA of the unbound complex
-nstruct 30 #number of desired number of ensembles
-relax:fast #Do 5 rounds of FastRelax
-out:pdb_gz #Compress (gzip) output pdbs
-out:prefix relax_ #Prefix for output pdb names
-out:path:all output #name of folder to store both pdb and score
```

   (c) Normal Mode Analysis

```
$ /path/to/Rosetta/main/source/bin/rosetta_scripts.linuxgccrelease
    @nma_flags
@nma_flags
-in:file:s complex_A.pdb #chainA of the unbound complex
-nstruct 30 #number of desired number of ensembles
-parser:protocol nma.xml #xml file for nma mover
-out:prefix nma_ #Prefix for output pdb names
-out:path:all output #name of the folder to store both pdb and score
```

```
nma.xml
<ROSETTASCRIPTS>
# This protocol mixes motion along the first 5 normal modes with a
    perturbation of 1.
    <SCOREFXNS>
            <ScoreFunction name="bn15_cart" weights="ref2015_cart"/>
    </SCOREFXNS>
    <RESIDUE_SELECTORS>
    </RESIDUE_SELECTORS>
    <TASKOPERATIONS>
    </TASKOPERATIONS>
    <FILTERS>
    </FILTERS>
    <MOVERS>
            <NormalModeRelax name="nma"
                    cartesian="true"
                    centroid="false"
                    scorefxn="bn15_cart"
                    nmodes="5"
                    mix_modes="true"
                    pertscale="1.0"
                    randomselect="false"
                    relaxmode="relax"
                    nsample="20"
                    cartesian_minimize="false"/>
    </MOVERS>
    <APPLY_TO_POSE>
    </APPLY_TO_POSE>
    <PROTOCOLS>
            <Add mover="nma"/>
    </PROTOCOLS>
    <OUTPUT scorefxn="bn15_cart"/>
</ROSETTASCRIPTS>
```

5. Prepacking

```
$ /path/to/Rosetta/main/source/bin/docking_prepack_protocol.linuxgccrelease
```

```
   @docking_prepack_flags

@docking_prepack_flags
-in:file:s complex.pdb #unbound complex with its components arbitrarily
   translated and rotated
-in:file:native native.pdb #native bound complex
-in:membrane #recognize membrane options
-mp:lipids:composition POPE #Type of lipids to use in implicit model
   representation.
-mp:lipids:temperature 35.0 #Temperature at which the lipid composition
   parameters were measured
-mp:lipids:has_pore 0 #use or not pore estimation
-score:weights 'franklin2023' #name of score function, options: franklin2019/
   franklin2023/mpframework_docking_fa_2015
-mp:setup:spanfiles complex.span #Spanning topology file of complex
-mp:setup:span1 chainA.span #Spanning topology file of partner1
-mp:setup:span2 chainB.span #Spanning topology file of partner2
-partners C_A #defines docking partners by ChainID, example: docking chains L+H
   with A is LH_A
-ensemble1 1ensemble.txt #Ensemble mode for partner1 on. 1ensemble.txt: path to
   partner1 ensembles
-ensemble2 2ensemble.txt #Ensemble mode for partner2 on. 2ensemble.txt: path to
   partner2 ensembles
-nstruct 1 #Number of prepacked structures of ensemble backbone
-ignore_zero_occupancy false #discard coordinate information for missing atoms
-ex1 #use extra chi1 sub-rotamers for all residues that pass the extrachi_cutoff
-ex2aro #use extra chi2 sub-rotamers for all aromatic residues
-out:suffix _ppk #Prefix for output pdb names
-overwrite #overwrite pdb files
```

6. Docking

```
$ /path/to/Rosetta/main/source/bin/mp_dock.linuxgccrelease @docking_flags
@docking_flags
-in:file:s complex.pdb #unbound complex with its components arbitrarily
   translated and rotated
-in:file:native native.pdb #native bound complex
-nstruct 5000 #number of docked complexes
-mp:setup:spanfiles complex.span #Spanning topology file of complex
-in:membrane #recognize membrane options
-mp:lipids:has_pore 0 #use or not pore estimation
-mp:lipids:composition POPE #Type of lipids to use in implicit model
   representation.
-mp:lipids:temperature 35.0 #Temperature at which the lipid composition
   parameters were measured
-dock_pert 3 8 #perturbations to each partner: 3A translation, 8degrees rotation
```

```
-ensemble1 1ensemble.txt.ensemble #Ensemble mode for partner1 on. 1ensemble.txt:
    path to partner1 ensembles
-ensemble2 2ensemble.txt.ensemble #Ensemble mode for partner2 on. 2ensemble.txt:
    path to partner2 ensembles
-partners C_A #defines docking partners by ChainID, example: docking chains L+H
   with A is LH_A
-docking_low_res_score motif_dock_score #Define low resolution docking score
   function, options: motif_dock_score,mpframework_docking_cen_2015(default
   option with -in:membrane flag)
-mh:path:scores_BB_BB /path to Rosetta/main/database/additional_protocol_data/
   motif_dock/xh_16_ #motif hash data for scoring, flag is only for
   motif_dock_score
-mh:score:use_ss1 false #flag is only for motif_dock_score
-mh:score:use_ss2 false #flag is only for motif_dock_score
-mh:score:use_aa1 true #flag is only for motif_dock_score
-mh:score:use_aa2 true #flag is only for motif_dock_score
-ignore_zero_occupancy false #discard coordinates information for missing atoms
-ex1 #use extra chi1 sub-rotamers for all residues that pass the extrachi_cutoff
-ex2aro #use extra chi2 sub-rotamers for all aromatic residues
-score:weights franklin2023 #score function for docking, options:franklin2019/
   franklin2023/mpframework_docking_fa_2015
-score:pack_weights franklin2023 #score function for packing, options:
   franklin2019/franklin2023/mpframework_docking_fa_2015)
-out:path:all highres_output #name of folder to store both pdb and score
-out:pdb_gz #Compress (gzip) output pdbs
-out:file:scorefile updated_docking_fa23.sc #score file for all output pdbs
-out:suffix _fa23 #prefix to putput pdb
-run:multiple_processes_writing_to_one_directory #when mpi is used all output
   files are written in one directory
```

7. Alphafold structure preparation

We predicted structures of complexes using the AlphaFold multimer using the protocols described in colabfold database. We selected rank0 pdb structures and used them as starting complexes by renaming them as complex.pdb. Steps 4-6 were repeated for rigid or ensemble docking protocols.

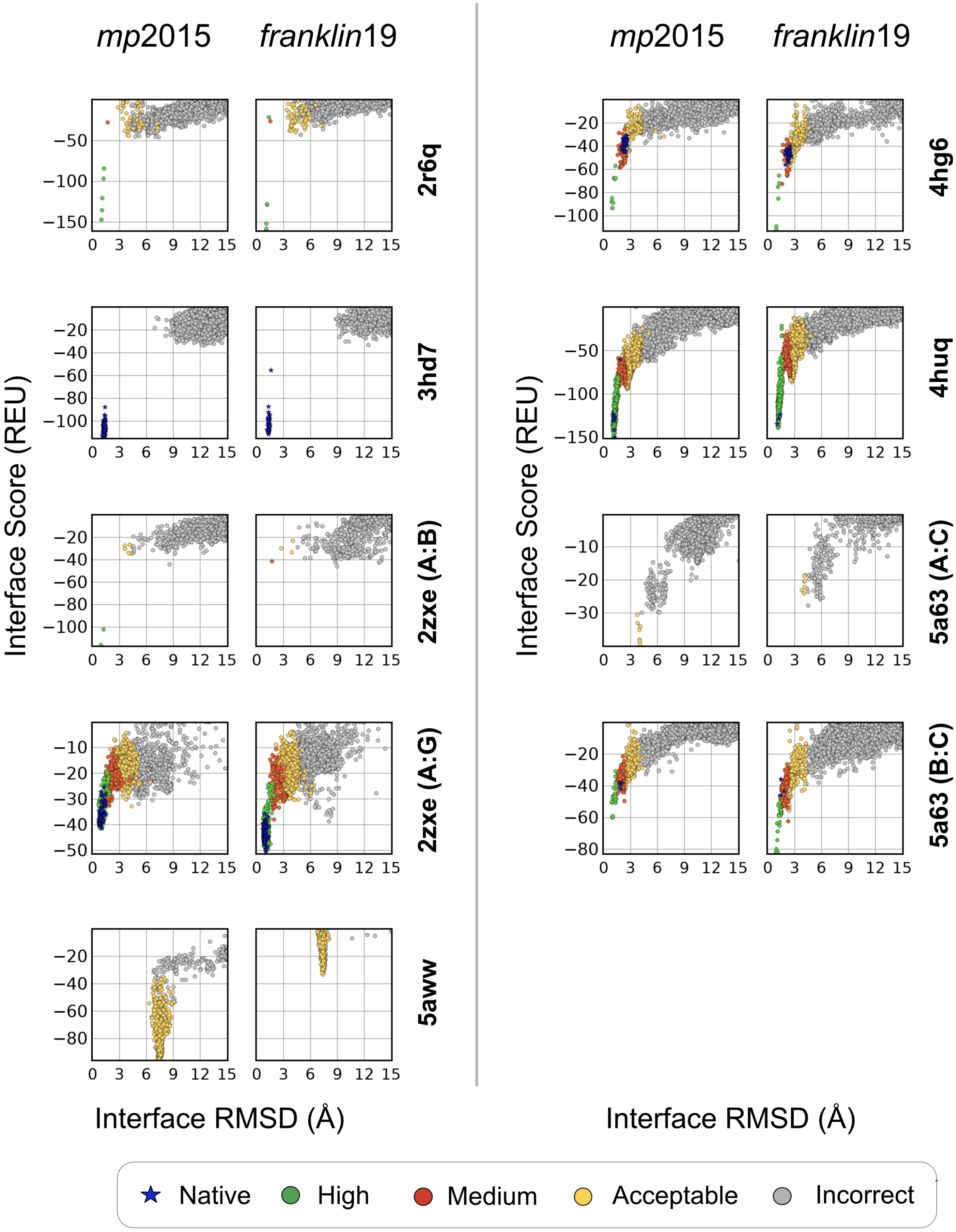


**Fig. S1. Rigid-body docking performance** for bound protein targets. Plots show the interface Score (REU) vs all-atom C$\alpha$ rmsd (Å). Blue points denote the refined native structures. (colors : green = high quality, red = moderate quality, yellow = acceptable quality, gray = incorrect)

#### References.

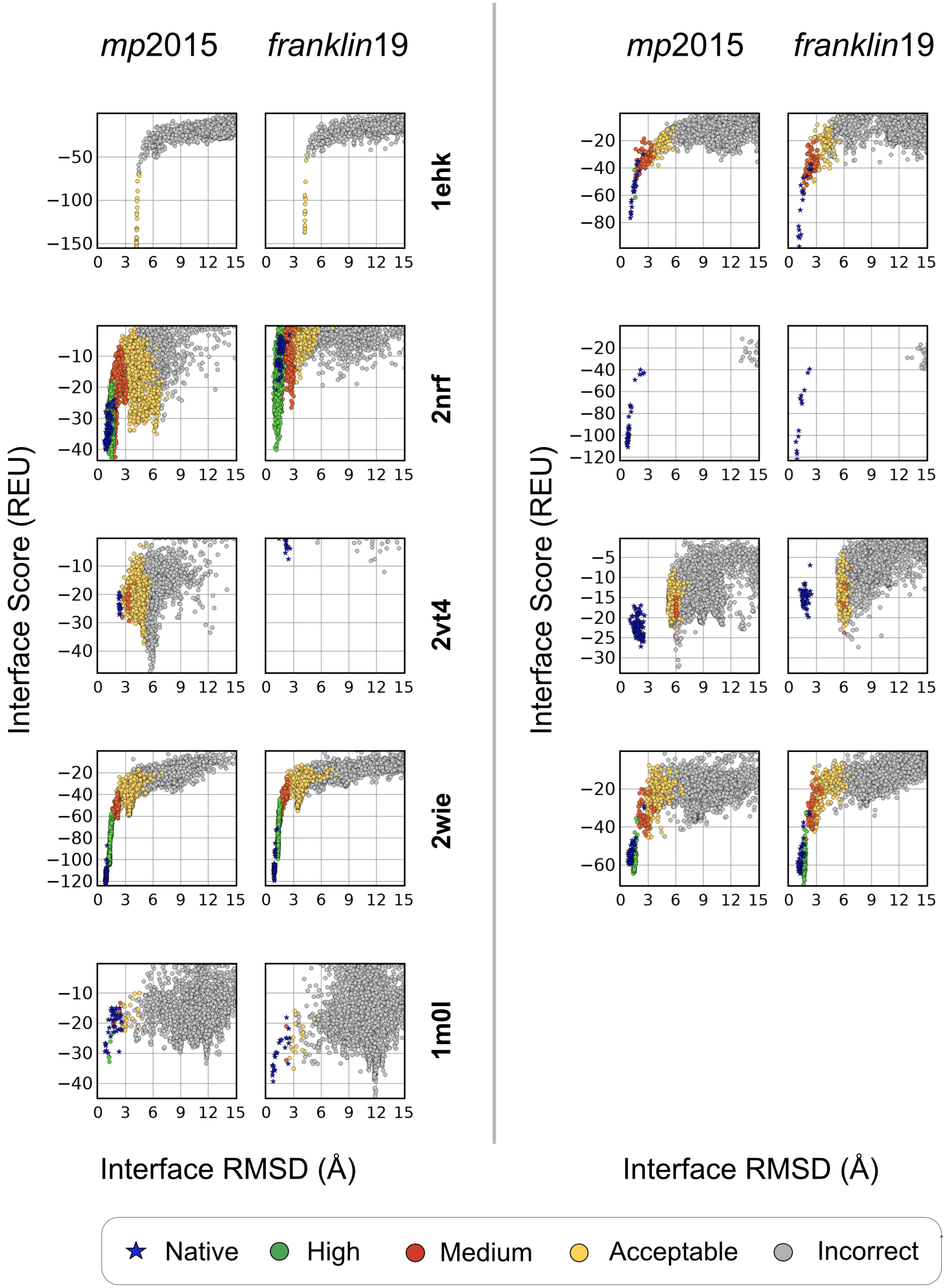


**Fig. S2. Rigid-body docking performance** for rigid protein targets where $RMSD_{BU} < 1.5$Å. Plots show the interface Score (REU) vs all-atom C$\alpha$ rmsd (Å). Blue points denote the refined native structures. (colors : green = high quality, red = moderate quality, yellow = acceptable quality, gray = incorrect)

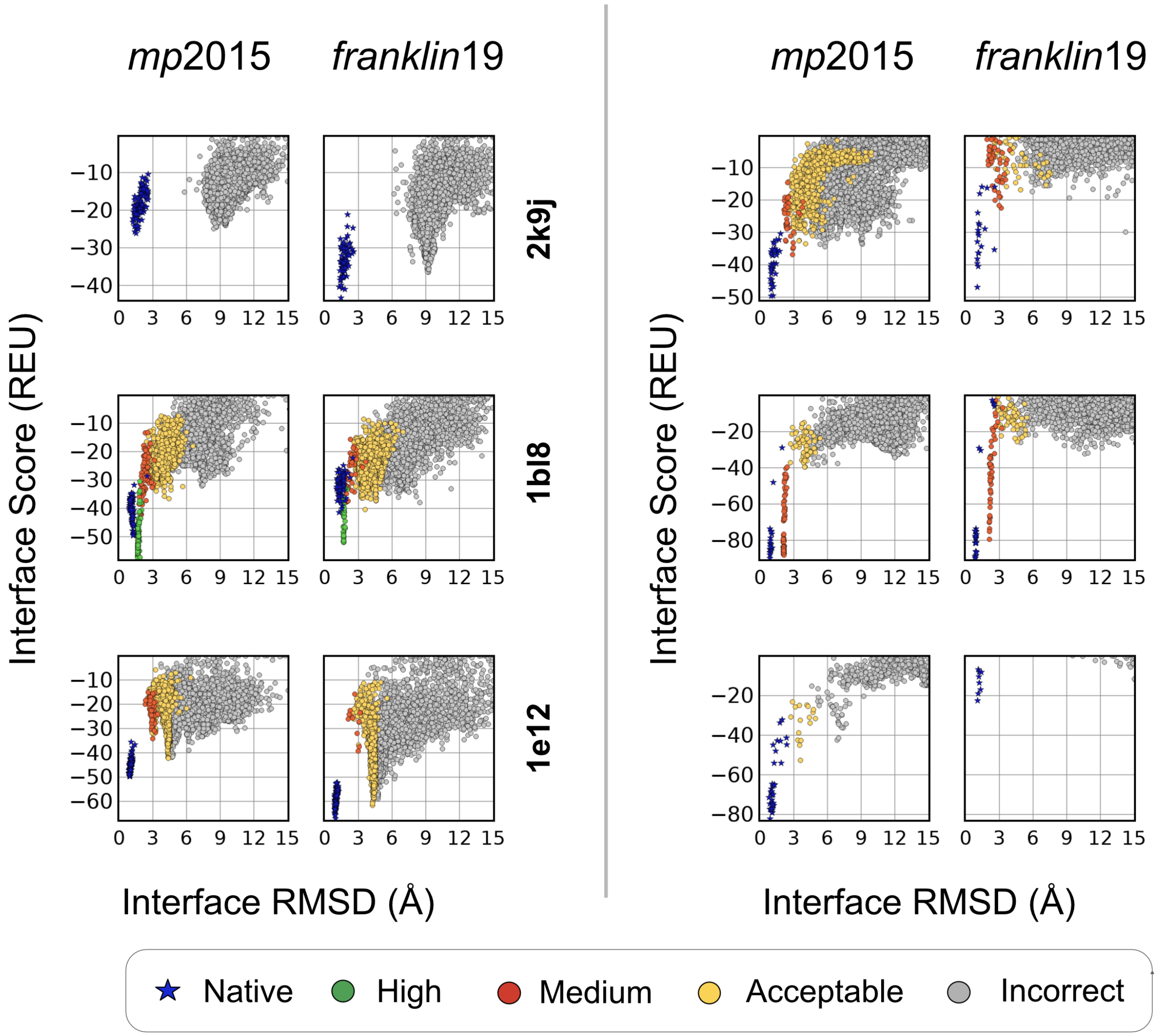


**Fig. S3. Rigid-body docking performance** for medium flexibility protein targets where $1.5\text{Å} < \text{RMSD}_{\text{BU}} < 2.5\text{Å}$. Plots show the interface Score (REU) vs all-atom C$\alpha$ rmsd (Å). Blue points denote the refined native structures. (colors : green = high quality, red = moderate quality, yellow = acceptable quality, gray = incorrect)

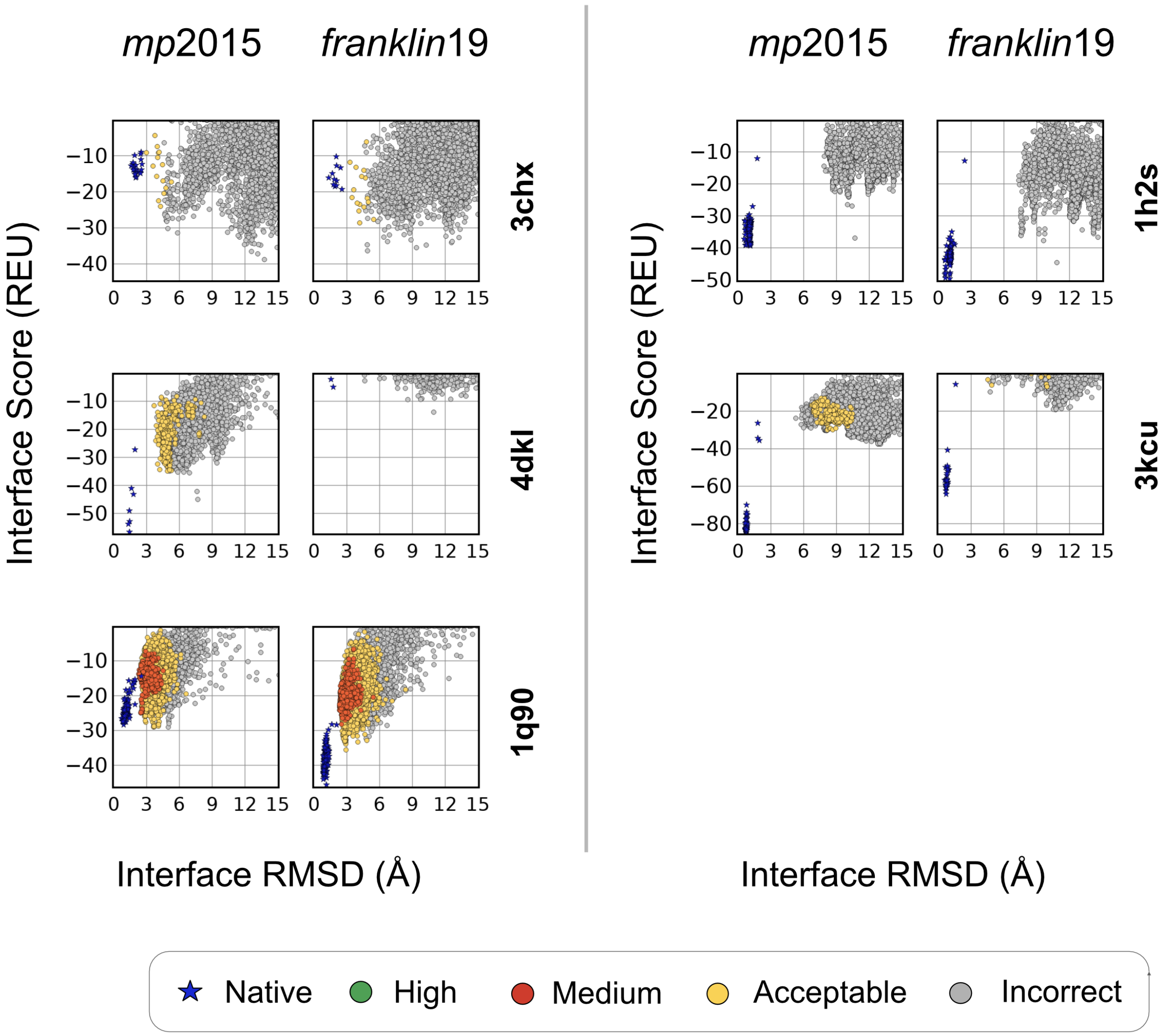


**Fig. S4. Rigid-body docking performance** for high flexibility protein targets where $\mathrm{RMSD_{BU}} > 2.5\text{Å}$. Plots show the interface Score (REU) vs all-atom C$\alpha$ rmsd (Å). Blue points denote the refined native structures. (colors : green = high quality, red = moderate quality, yellow = acceptable quality, gray = incorrect)

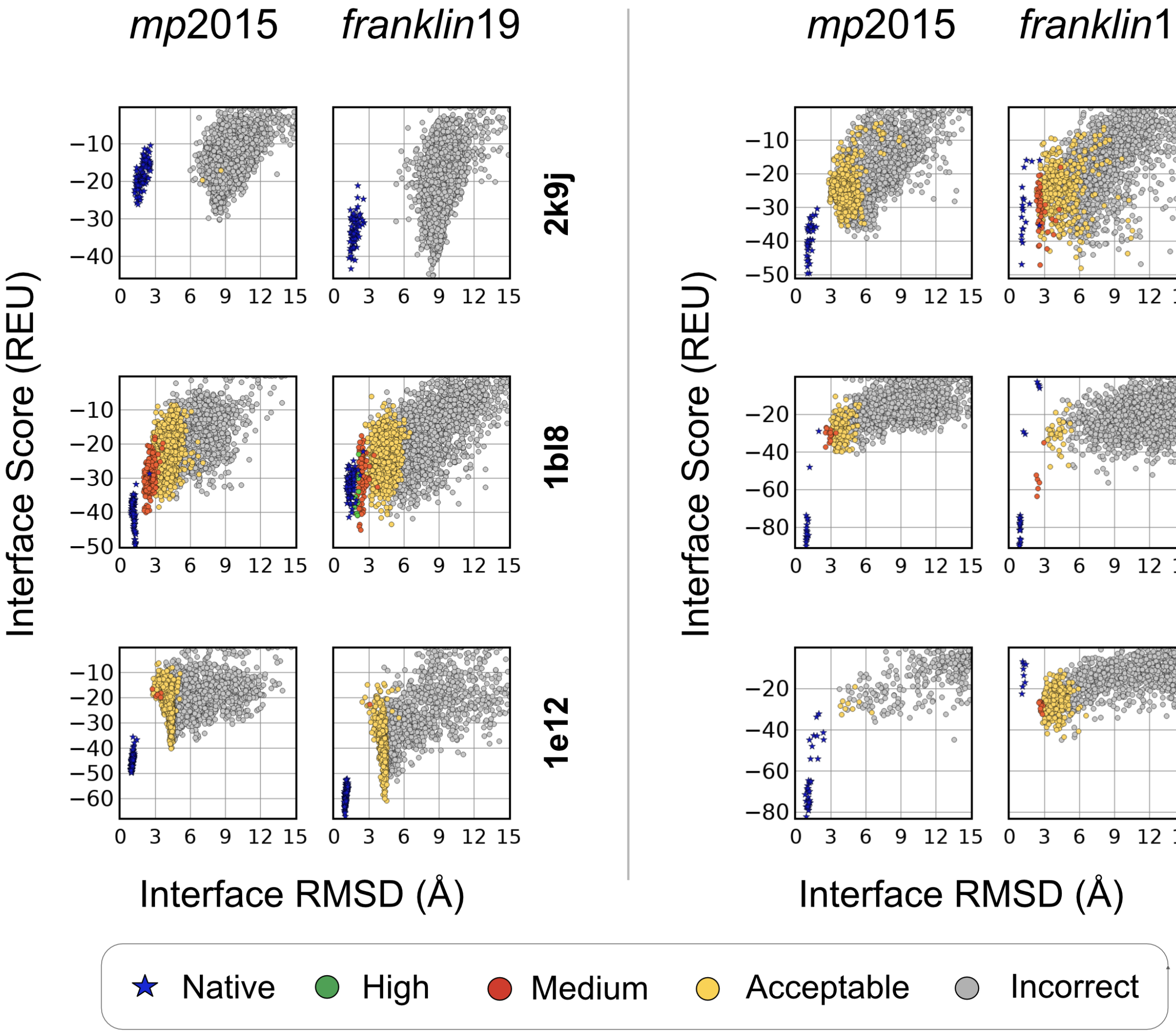


**Fig. S5. Ensemble-MPdock performance** for medium flexibility protein targets where $1.5\text{Å} < \text{RMSD}_{\text{BU}} < 2.5\text{Å}$. Plots show the interface Score (REU) vs all-atom C$\alpha$ rmsd (Å). Blue points denote the refined native structures. (colors : green = high quality, red = moderate quality, yellow = acceptable quality, gray = incorrect)

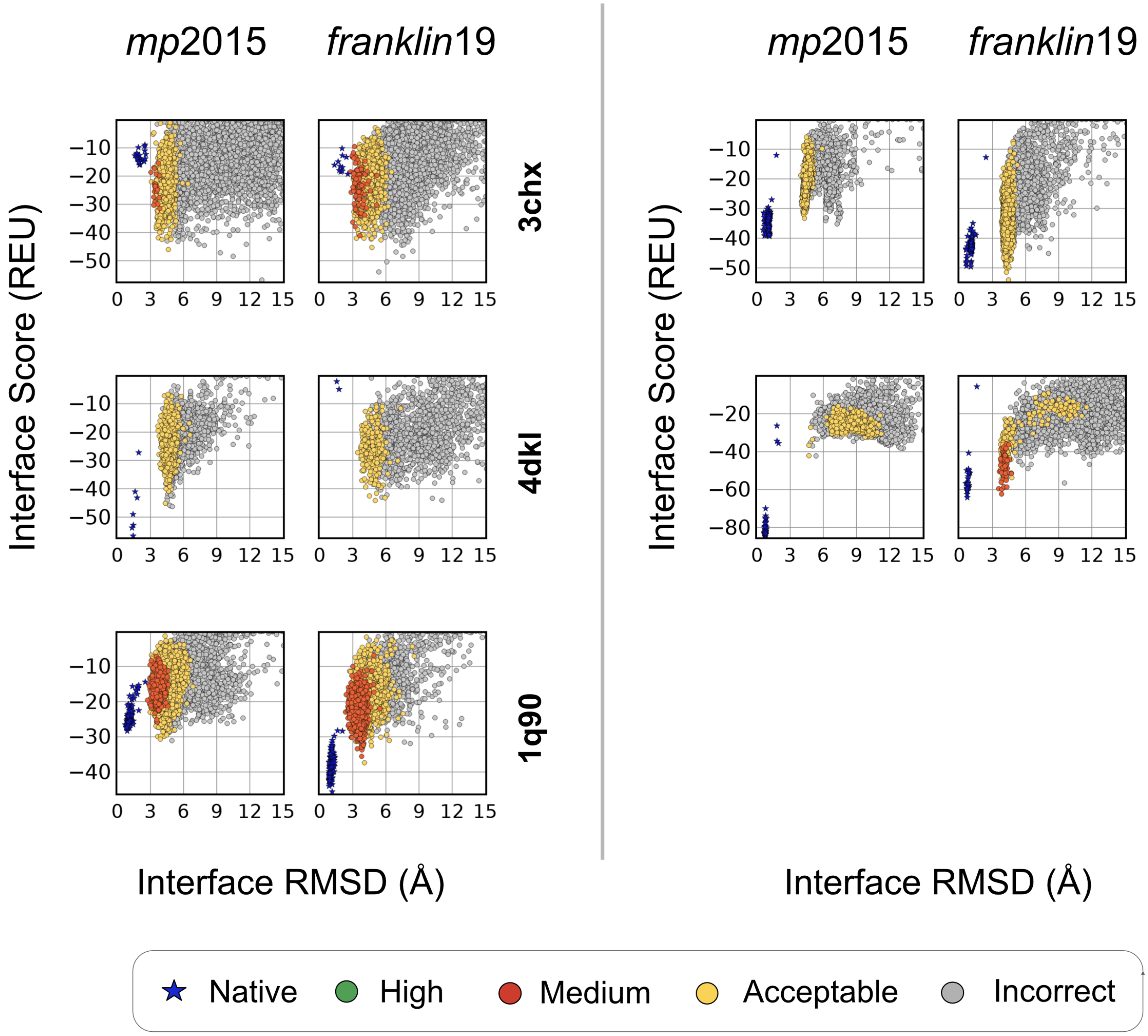


**Fig. S6. Ensemble-MPDock performance** for high flexibility protein targets where $\text{RMSD}_{\text{BU}} > 2.5\text{Å}$. Plots show the interface Score (REU) vs all-atom C$\alpha$ rmsd (Å). Blue points denote the refined native structures. (colors : green = high quality, red = moderate quality, yellow = acceptable quality, gray = incorrect)

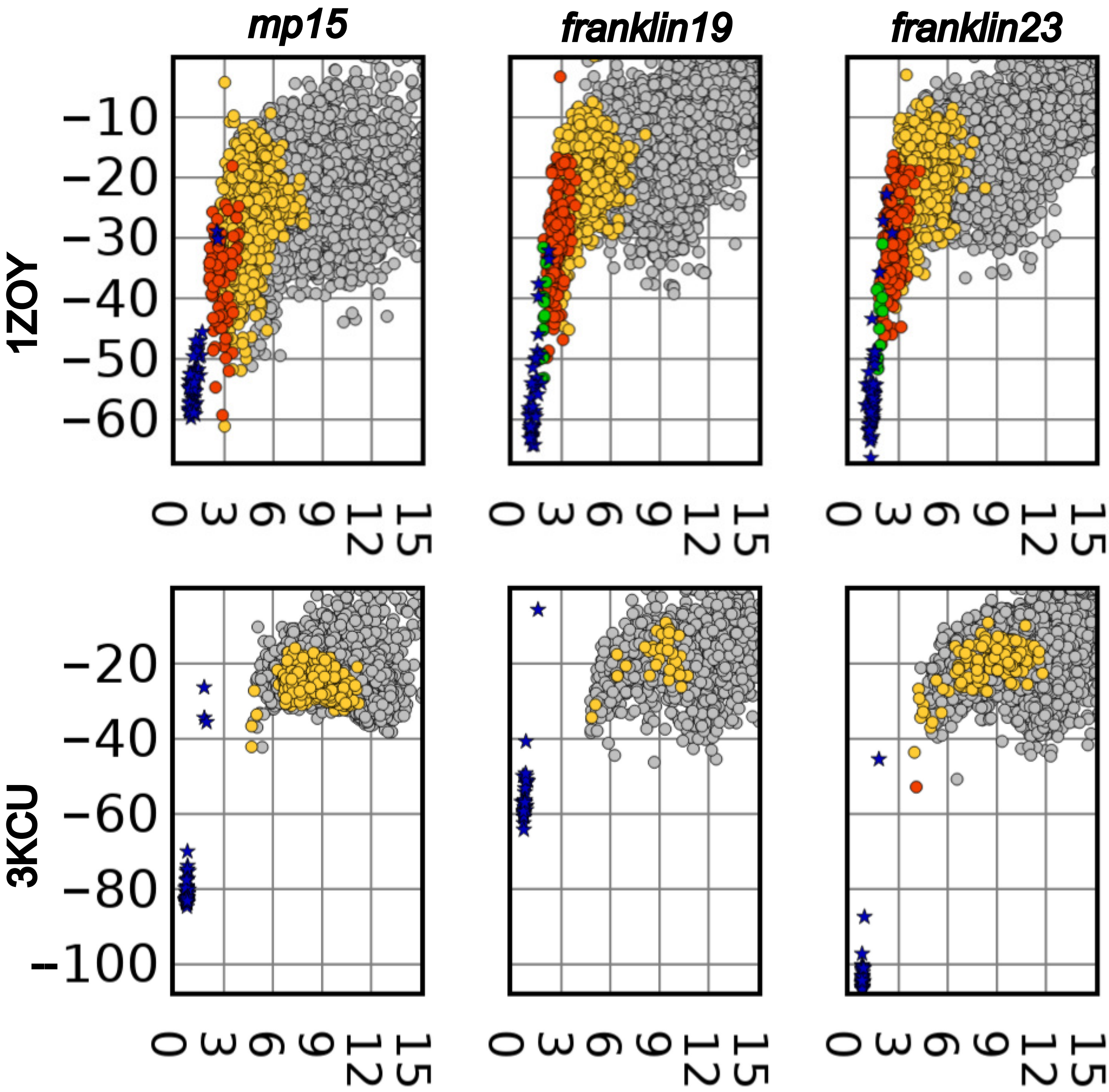


**Fig. S7. Ensemble-MPDock performance** for protein targets 1ZOY (mitochondrial respiratory complex II, $\mathrm{RMSD}_{BU}$ = 1.20 Å) and 3KCU (Portable formate transporter, $\mathrm{RMSD}_{BU}$ = 3.56 Å). Plots show the interface Score (REU) vs all-atom $C\alpha$ rmsd (Å). Blue points denote the refined native structures. (colors : green = high quality, red = moderate quality, yellow = acceptable quality, gray = incorrect)

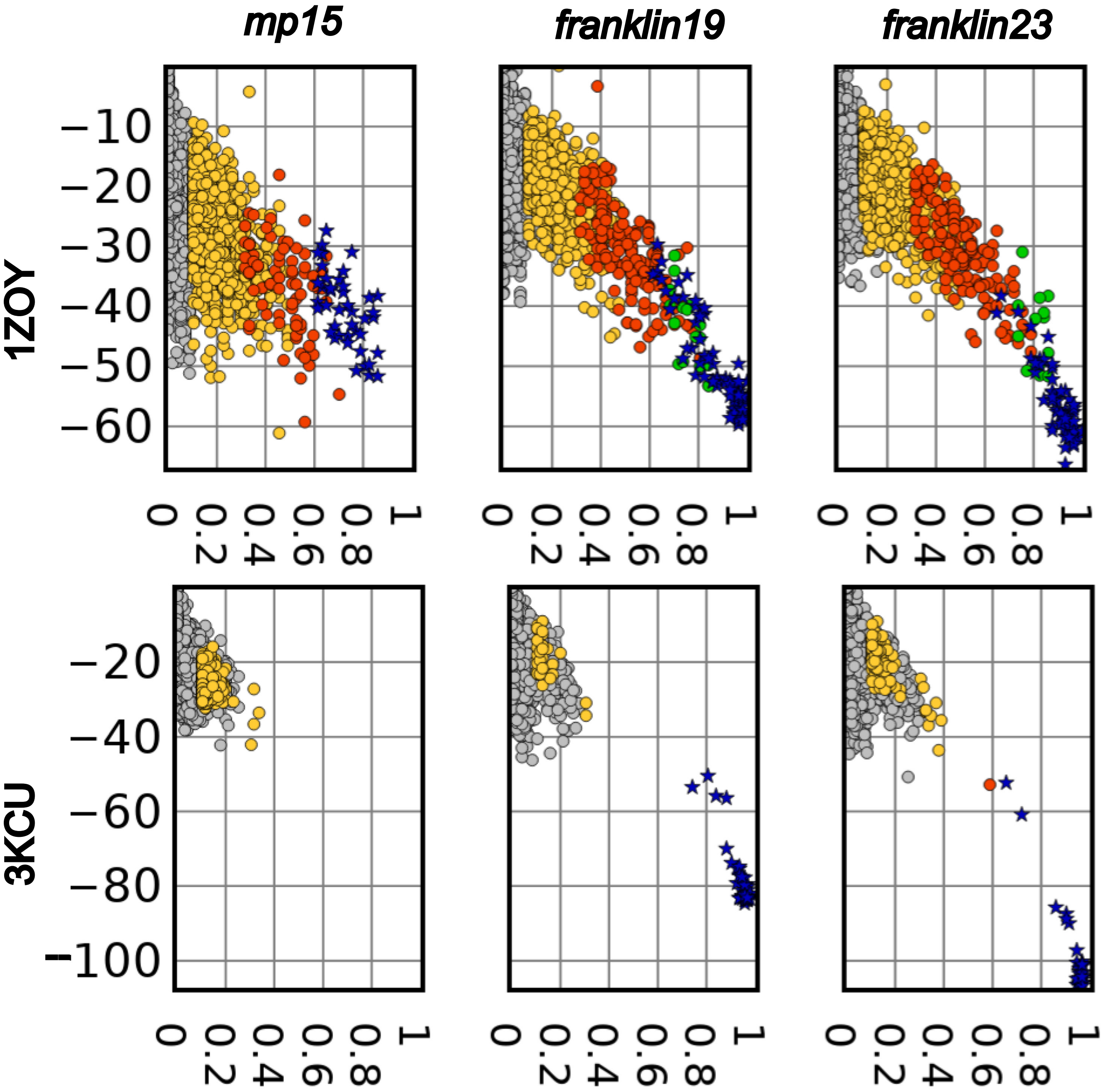


**Fig. S8. Ensemble-MPDock performance** for protein targets 1ZOY (mitochondrial respiratory complex II, $RMSD_{BU}$ = 1.20 Å) and 3KCU (Portable formate transporter, $RMSD_{BU}$ = 3.56 Å). Plots show the fraction of native-like contacts vs all-atom C$\alpha$ rmsd (Å). Blue points denote the refined native structures. (colors : green = high quality, red = moderate quality, yellow = acceptable quality, gray = incorrect)

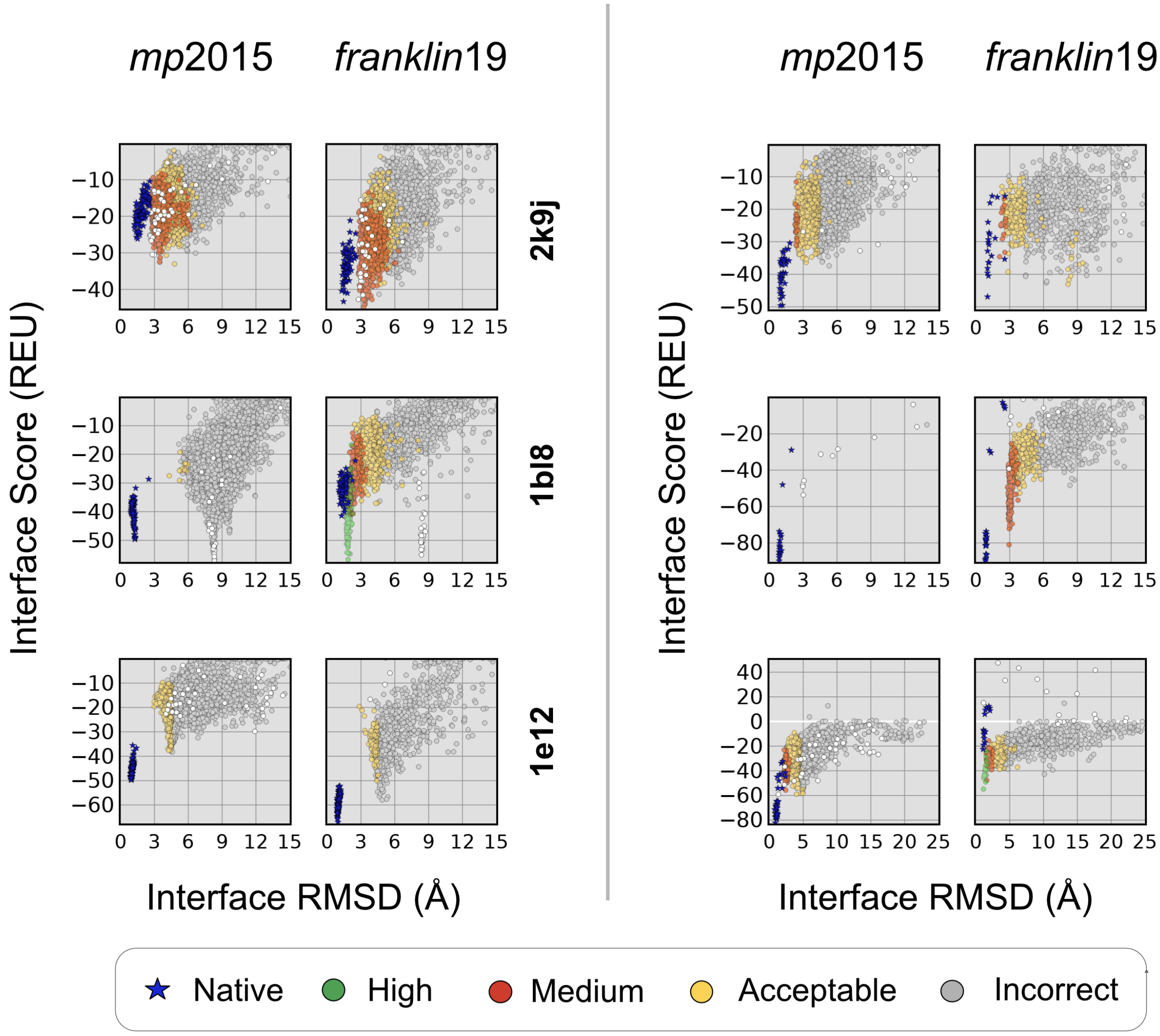


**Fig. S9. Performance of MPDock with Alphafold-2 predicted structures** for medium flexibility protein targets where $1.5\text{Å} < \text{RMSD}_{\text{BU}} < 2.5\text{Å}$. Plots show the interface Score (REU) vs all-atom C$\alpha$ rmsd (Å). Blue points denote the refined native structures. (colors : green = high quality, red = moderate quality, yellow = acceptable quality, gray = incorrect)

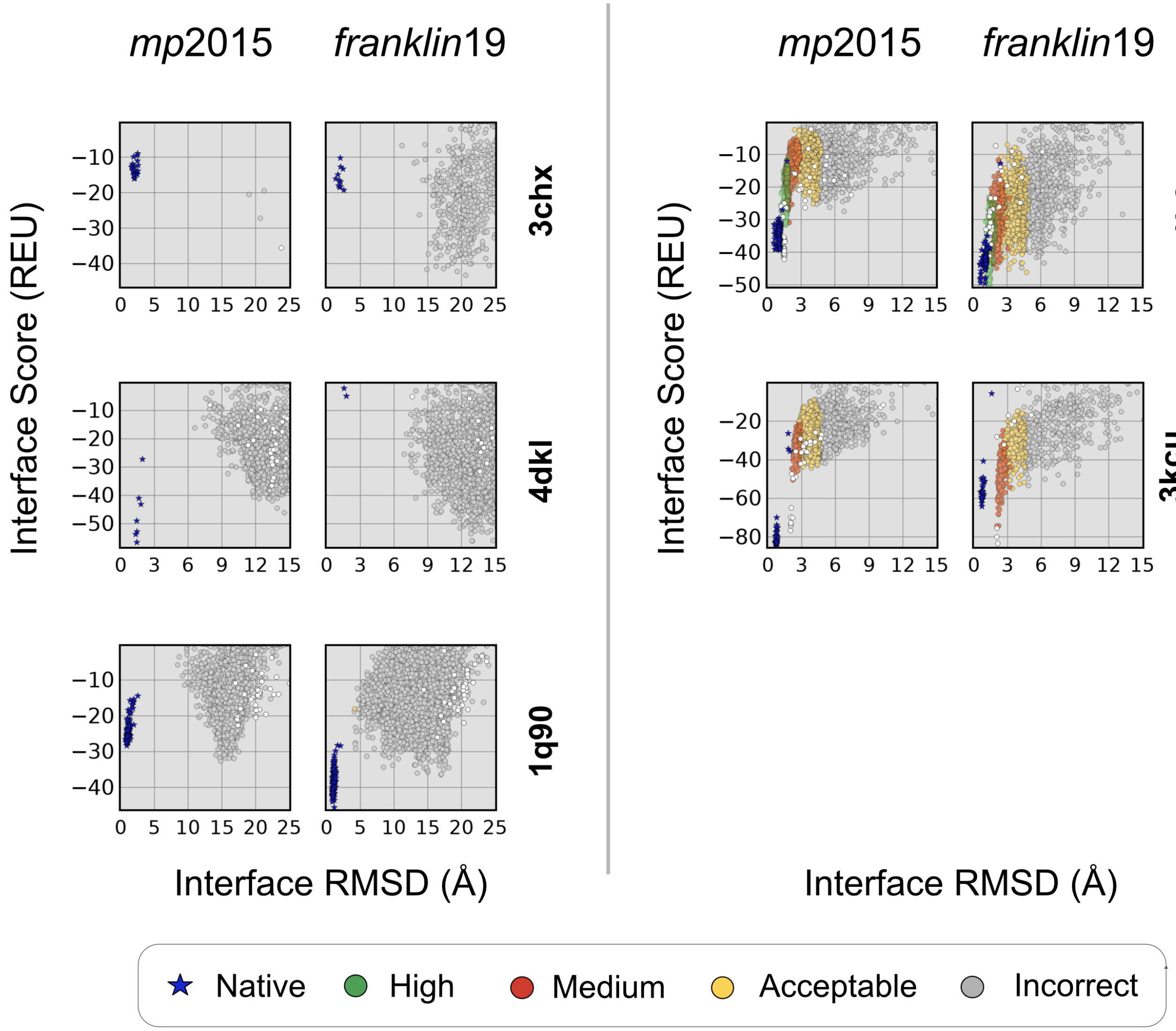


**Fig. S10. Performance of MPDock with Alphafold-2 predicted structures** for high flexibility protein targets where $RMSD_{BU} > 2.5Å$. Plots show the interface Score (REU) vs all-atom C$\alpha$ rmsd (Å). Blue points denote the refined native structures. (colors : green = high quality, red = moderate quality, yellow = acceptable quality, gray = incorrect)
